# Stabilizing gas in aqueous liquids with metal–organic frameworks for persistent acoustic transduction

**DOI:** 10.64898/2026.10.27.762357

**Authors:** Wenliang Wang, Weilong He, Miranda V Walter, Zixiang Tong, Xiangping Liu, Binita Shrestha, Brinkley Artman, Ju-Chun Hsieh, Sihan Yu, Jinmo Jeong, Ilya Pyatnitskiy, Sihan Yu, Mengmeng Yao, Haiyue Xu, Jacob Baker, Lehu Bu, Yujie Zhang, Noah B. Stern, Bing-Ze Lin, Matthew R. Lowerison, Pengfei Song, Tyrone Porter, Jarad Mason, Jin Yang, Huiliang Wang

## Abstract

Gas is the most efficient mediator of acoustic energy owing to its high compressibility, yet its thermodynamic instability in aqueous environments makes it inherently transient, limiting sustained ultrasound–biological interfacing. Here, we show that gas can be stabilized, confined, and manipulated in the liquid phase to function as a persistent acoustic transducer. By trapping gas within designed microporous metal–organic framework (MOF) nanocrystals, interfacial forces establish a stable ternary-phase equilibrium that suppresses water intrusion and prevents gas collapse, enabling reversible, non-destructive oscillations under ultrasound stimulation and providing design principles for stabilizing gas in liquid environments. By leveraging high-resolution ultra-high-speed digital image correlation (DIC), we unveil the previously inaccessible spatiotemporal dynamics of confined gas oscillations within these nanotransducers under ultrasound stimulation, revealing localized amplification of the surrounding acoustic field and establishing a quantitative structure–function relationship for programmable acoustic transduction. These aqueous-phase gas nanotransducers exhibit sustained acoustic amplification over millions of excitation cycles and remain functional after prolonged storage (>2 years) under physiological conditions. *In vitro* and *in vivo*, this platform enables long-term ultrasound imaging and sustained, non-invasive neuromodulation by converting confined gas oscillations into persistent and amplified mechanical actuation. Our findings establish a generalizable design framework for stabilizing gas in liquid environments to achieve persistent and programmable acoustic transduction, thereby enabling sustained ultrasound–biological interfacing and opening opportunities for gas-based therapeutics.

## Introduction

The ability to transduce energy between physical and biological systems underpins technologies for sensing, actuation, and communication.(*1–3*) In acoustic systems, gas is the most efficient mediator of energy conversion owing to its high compressibility, enabling efficient interactions between acoustic waves and biological systems.(*4–6*) Yet the same behavior that makes gas acoustically powerful also makes it difficult to use as a persistent transducer in aqueous biological environments as gases dissolve, diffuse, or collapse under pressure.(*1*, *7*) This intrinsic instability has limited the development of long-lived, controllable ultrasound-based technologies for deep-tissue interfacing.

Clinically used microbubbles have established gas as a powerful acoustic mediator for ultrasound imaging and therapy, yet their micrometre size, intravascular confinement, limited circulation lifetime, and susceptibility to acoustic destruction restrict their use in sustained ultrasound–biological interfacing.(*1*, *7*, *8*) Nanoscale gas systems, including nanobubbles and gas vesicles (GVs) offer improved nanoscale stability and acoustic performance, and have enabled ultrasound-based visualization and control of cellular processes.(*2*, *6*, *9*) However, gas in these systems remains prone to dissolution, leakage, or structural collapse under repeated acoustic excitation, preventing persistent and programmable acoustic transduction in biological environments.(*7*, *10*, *11*) Furthermore, their dependence on cold-chain storage imposes significant logistical burdens, further hindering widespread clinical adoption.(*12*, *13*) Thus, a central unresolved question is whether gas—an inherently unstable phase in liquid—can be stabilized and confined to enable persistent, programmable acoustic transduction, a capability that existing systems lack.

Recently, porous materials have been used to stabilize gas molecules in liquids, resulting in a new class of fluids known as porous liquids.(*14*) In these systems, solvent molecules are excluded from the pores of the dispersed material through steric or thermodynamic effects.(*15–17*) In particular, microporous materials can be designed to adsorb high densities of gases in aqueous environments by combining a hydrophobic internal pore surface with a hydrophilic external particle surface to form fluids termed “microporous water.”(*18*) Though the increased gas carrying capacities of microporous water have been leveraged for aqueous-phase gas transport, the acoustical properties of these systems, which may be of interest for ultrasound applications, have yet to be investigated. As such, we hypothesized that microporous water might provide new opportunities for preventing interfacial destabilization of gas confined in liquids under dynamic acoustic excitation, as well as serving as a platform for establishing the largely unexplored structure–function relationships that govern gas retention, confined oscillation, and long-term acoustic durability.

Here, we establish a generalizable physical framework and rational design strategy to stabilize and program gas-mediated acoustic transduction in aqueous environments. We show that gas can be confined within hydrophobic porous MOFs nanocrystals, where interfacial forces govern a three-phase equilibrium that prevents collapse while enabling reversible, non-inertial oscillations under ultrasound. Guided by this principle, we systematically engineer MOFs as liquid-phase gas systems. These systems exhibit exceptional stability, retaining gas for over two years under ambient conditions and remaining functional after prolonged incubation at elevated temperatures. Under acoustic excitation, confined gas domains undergo restricted oscillations that generate persistent and amplified acoustic responses over millions of cycles (**Supplementary Video 1**), far exceeding the durability of existing agents. By integrating ultra-high-speed digital image correlation (DIC), a full-field deformation measurement technique, we directly visualize and quantify localized acoustic field amplification, revealing structure–function relationships that guide programmable design. Furthermore, *in vitro* and *in vivo*, these nanotransducers enable long-term ultrasound imaging and durable targeted neuromodulation, establishing a new class of non-collapsing acoustic transducers for persistent deep-tissue ultrasound–biological interfacing.

### Designing MOFs as liquid-phase gas nanotransducers

To develop a method for identifying porous materials that may serve as promising candidates for gas stabilization in aqueous environments, we first considered the physical principles governing gas confinement in the liquid phase, drawing on prior work.(*15*, *17*) In natural systems, the diving bell spider (*Argyroneta aquatica*) maintains a stable air layer underwater by trapping gas at hydrophobic interfaces (**Fig. 1a**), illustrating how interfacial forces can counteract hydrostatic pressure and stabilize gas in liquid. (*19*) A physical model governed by Laplace’s law was developed to describe the three-phase equilibrium of confined gas within porous structures (**Fig. 1b–d and Supplementary Information, Equations S1–S5**). Following the principle (**Fig. 1c,d**; see details in **Supplementary Information**), we describe the interfacial balance of forces governing gas confinement within nanoporous structures. Gas bubbles trapped inside pores of porous materials are subjected to inward hydrostatic pressure (*P*_w_), outward gas pressure (*P*_a_), and surface tension (*γ*). The direction of surface tension depends on the wetting properties of the pore walls. For hydrophobic porous materials (*θ* > 90°, cos *θ* < 0), surface tension acts outward, counteracting hydrostatic pressure and preventing water intrusion, thereby stabilizing gas pockets within the pore channels. Conversely, for hydrophilic porous materials (0 < *θ* < 90°, cos *θ* > 0), surface tension acts inward, promoting water penetration into the pores. Beyond surface wettability, pore architecture is also expected to modulate the interfacial force balance (**Fig. 1d**), enabling programmable control over the critical pressure for water intrusion, above which confined gas becomes destabilized and expelled from the nanopores (see **Supplementary Information**). (*20*– *22*)

**Fig. 1.**
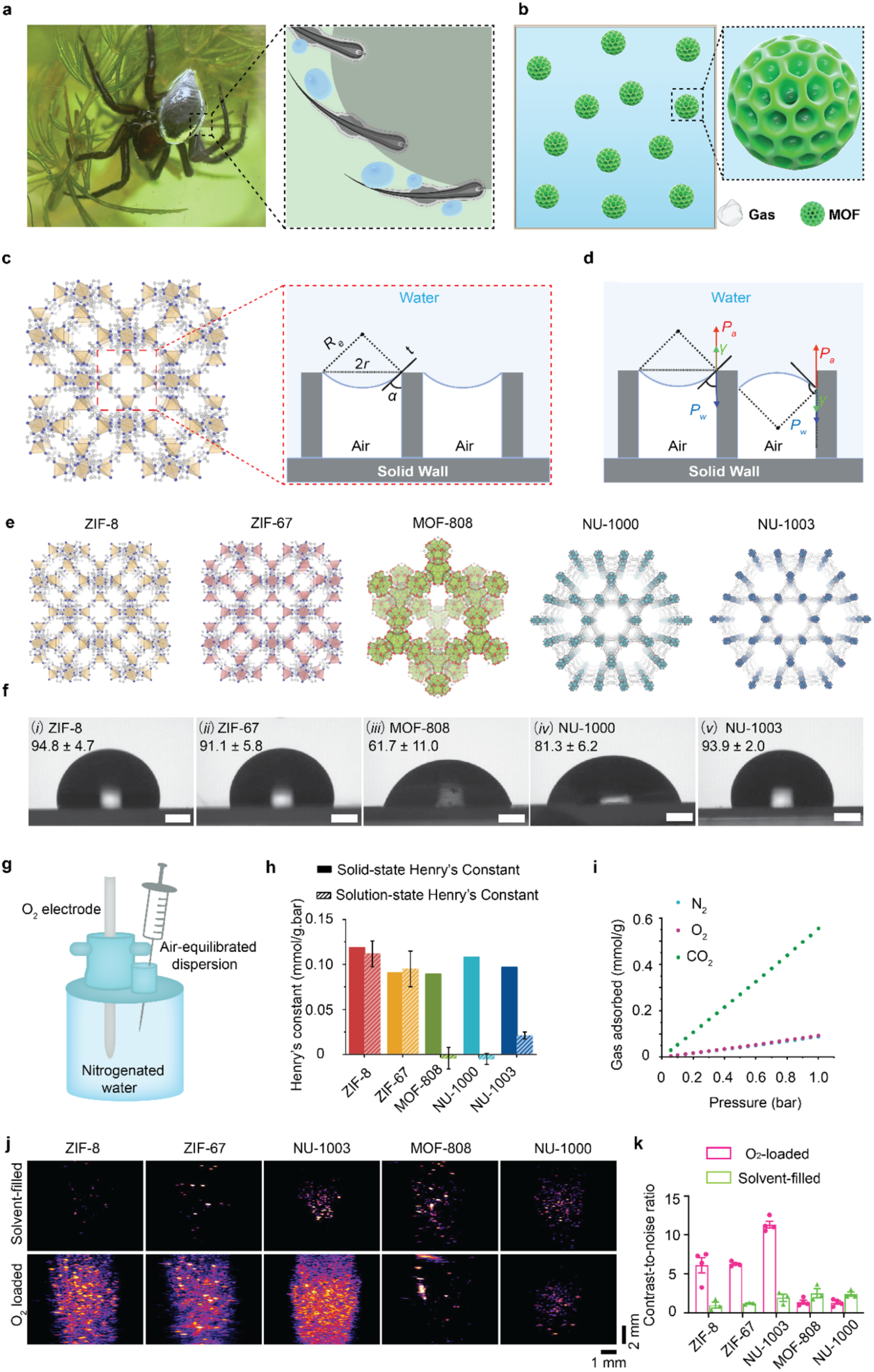
Microporous MOFs act as aqueous-phase gas nanotransducers. (a) Schematic illustration of the water spider, *Argyroneta aquatica*, which survives underwater by trapping air with its hydrophobic abdominal hairs. (b) Microporous metal–organic frameworks (MOFs) similarly confine gas within hydrophobic pore channels, maintaining stability in aqueous environments. (c) Schematic representation of gas confinement in hydrophobic pore channels. Side view of the air–water interface between two three-phase contact points (grey bars), separated by a distance of 2*r*. *R_e_* denotes the radius of curvature of the interface; *t* is the tangent to the curved interface at a contact point. The contact angle *θ* (measured across the liquid phase) is defined as *θ* = 180° − *α*, shown for *θ* > 90°. (d) Schematic illustration of the force balance governing gas confined within MOF pores with different surface properties. The system is subject to inward hydrostatic pressure (P_w_), outward gas pressure (P_a_), and surface tension (*γ*), the direction of which depends on the interfacial properties of the MOF inner surface. (e) Schematic representation of porous MOFs. (f) Contact-angle measurements of MOF pellets (n = 5), Scale bar: 0.5 mm. (g) Diagram of O_2_ release measurements used to calculate solution-state Henry’s constants for all microporous dispersions. (h) Henry’s constants for O_2_ adsorption in MOFs dispersed in water, compared with solid-state Henry’s constants calculated from O_2_ isotherms at 20 °C. All tested solutions were at a concentration of 10 mg/mL. (i) Solid-state CO_2_, N_2_, and O_2_ adsorption isotherms at 37 °C for ZIF-8. (j) Ultrasound imaging (27 MHz) of MOFs with or without O_2_ loading in aqueous solution at a concentration of 2 mg/mL. (k) Corresponding contrast-to-noise ratio (CNR) analysis.

To test this principle, we employed metal–organic framework (MOF) nanocrystals as a model system owing to their precisely tunable pore structures and surface chemistries (**Fig. 1e**). We synthesized a representative panel of MOFs, including ZIF-8, ZIF-67, MOF-808, NU-1000, and NU-1003 (**Fig. 1e**), among which ZIF-8 and ZIF-67 have previously been shown to retain dry micropores in aqueous environments.(*18*) Powder X-ray diffraction (XRD) and Brunauer– Emmett–Teller (BET) analyses confirmed their crystalline microporous structures (**Supplementary Figs. 1, 2**), while dynamic light scattering (DLS) and transmission electron microscopy (TEM) verified nanoscale uniformity (**Supplementary Fig. 3**). Next, we systematically investigated the surface properties of these MOFs. Contact angle (*θ*) measurements show that ZIF-8, ZIF-67, and NU-1003 exhibit hydrophobic surfaces, with values of 94.8°, 91.1°, and 93.9°, respectively (**Fig. 1e**). In contrast, MOF-808 and NU-1000 display hydrophilic behavior, with contact angles below 90° (**Fig. 1e)**. These results align with previous reports of hydrophobicity in ZIF-8 and ZIF-67 and indicate that NU-1003 might also retain porosity in water.(*18*) Thus, we next assessed the gas capture performance of these MOFs in aqueous environments.

Gas capacities for MOF nanocrystals in solution were determined by using an O_2_ electrode to measure the amount of O_2_ released from air-equilibrated samples into deoxygenated water. This allows the O_2_ adsorption capacity of the dispersed particles to be calculated and compared to its solid-state value (**Fig. 1g**). ZIF-8 and ZIF-67 dispersed in water exhibit gas sorption properties comparable to their solid-state counterparts as previously shown, whereas the gas capacities of MOF-808 and NU-1000 are effectively eliminated in solution (**Fig. 1h**).(*18*) Notably, NU-1003 retains a Henry’s constant in solution that is ∼22% of its solid-state value, indicating partial water intrusion and at least some retention of porosity. Structurally, NU-1003 contains two types of pore channels connected by ∼12 × 13 Å windows: large hexagonal channels (∼45 Å) and smaller triangular channels (∼17 Å) (**Supplementary Fig. 4**).(*23*) The triangular channels account for ∼31% of the total pore volume (0.2 cm³ g⁻¹ versus 0.45 cm³ g⁻¹ for hexagonal channels), potentially suggesting that water only intrudes into the large pore channels while the small pore channels remain dry and capable of adsorbing gas molecules. We further determined that different gases (non-polar gases, including CO_2_, N_2_, and O_2_) could be taken up by the MOF at a given pressure and stably stored at different temperatures (**Fig.1i** and **Supplementary Figs. 5, 6**). These findings indicate that contact angle measurements might be useful as a quick screening tool for evaluating water intrusion in MOFs, as confirmed by the full porosity exhibited by ZIF-8 and ZIF-67 and the partial porosity exhibited by NU-1003 through aqueous-phase oxygen-release measurements. Because these materials can confine and stabilize gas within aqueous environments, we proceeded to evaluate their effectiveness as potential acoustic energy mediators.

As anticipated, ZIF-8, ZIF-67, and NU-1003 all function as liquid-phase gas nanotransducers, exhibiting distinct acoustic backscattering behaviours under ultrasound excitation. Specifically, in a silicone rubber phantom evaluated with a 29 MHz scanning system, all three MOFs show pronounced enhancement of backscattering signals following oxygen loading, whereas only weak and sporadic signals are observed in the corresponding gas-free controls (**Fig. 1j,k**). In contrast, MOF-808 and NU-1000 nanocrystals exhibit negligible scattering signals regardless of gas loading (**Fig. 1j,k**), consistent with their limited gas adsorption capacity in aqueous environments (**Fig. 1h**). We further evaluated the imaging performance of ZIF-8 nanotransducers loaded with different gases, including CO₂, N₂, O₂, and air (**Supplementary Fig. 7**), which exhibited high contrast regardless of whether the MOF dispersions were initially equilibrated with air, N₂, or CO₂ (**Fig. 1i**). Together, these results demonstrate that liquid-phase gas nanotransducers can serve not only as ultrasound imaging reporters to enhance imaging contrast and resolution, but also as indicators of gas loading states within the nanostructures. However, the kinetic behaviour of confined gas under ultrasound remains poorly understood, and elucidating these dynamics is central to achieving programmable gas-state control and precise ultrasound–biological interfacing.

### Confined gas oscillations define a non-collapsing acoustic regime

We hypothesize that enhanced acoustic coupling arises from confined gas oscillations within MOF pore channels under ultrasound excitation, defining a non-collapsing oscillatory regime distinct from conventional cavitation. In this regime, gas domains remain spatially confined and undergo reversible oscillations without structural failure, enabling sustained and tunable acoustic responses. To test this hypothesis, we employ ultra-high-speed DIC to directly resolve transient acoustic stress fields generated by these nanotransducers.(*24*, *25*) MOF particles embedded in a gelatin hydrogel are subjected to laser-induced cavitation, generating controlled mechanical waves that mimic ultrasound propagation in biological tissues (**Fig. 2a,b**, **Supplementary Fig. 8 and Equations S7–S9**), where the underlying physical framework is detailed in **Supplementary Information.**(*26*, *27*) The resulting primary cavitation bubble produces a strong outward-propagating mechanical wave, which dynamically interacts with nearby MOF particles and induces secondary oscillations through bubble–particle coupling.(*28–30*) The temporal evolution of both the primary cavitation bubble and MOF clusters is captured using an ultra-high-speed camera at 1–10 million frames per second, enabling quantitative analysis of the resulting dynamic stress fields (**Fig. 2c**).

**Fig. 2.**
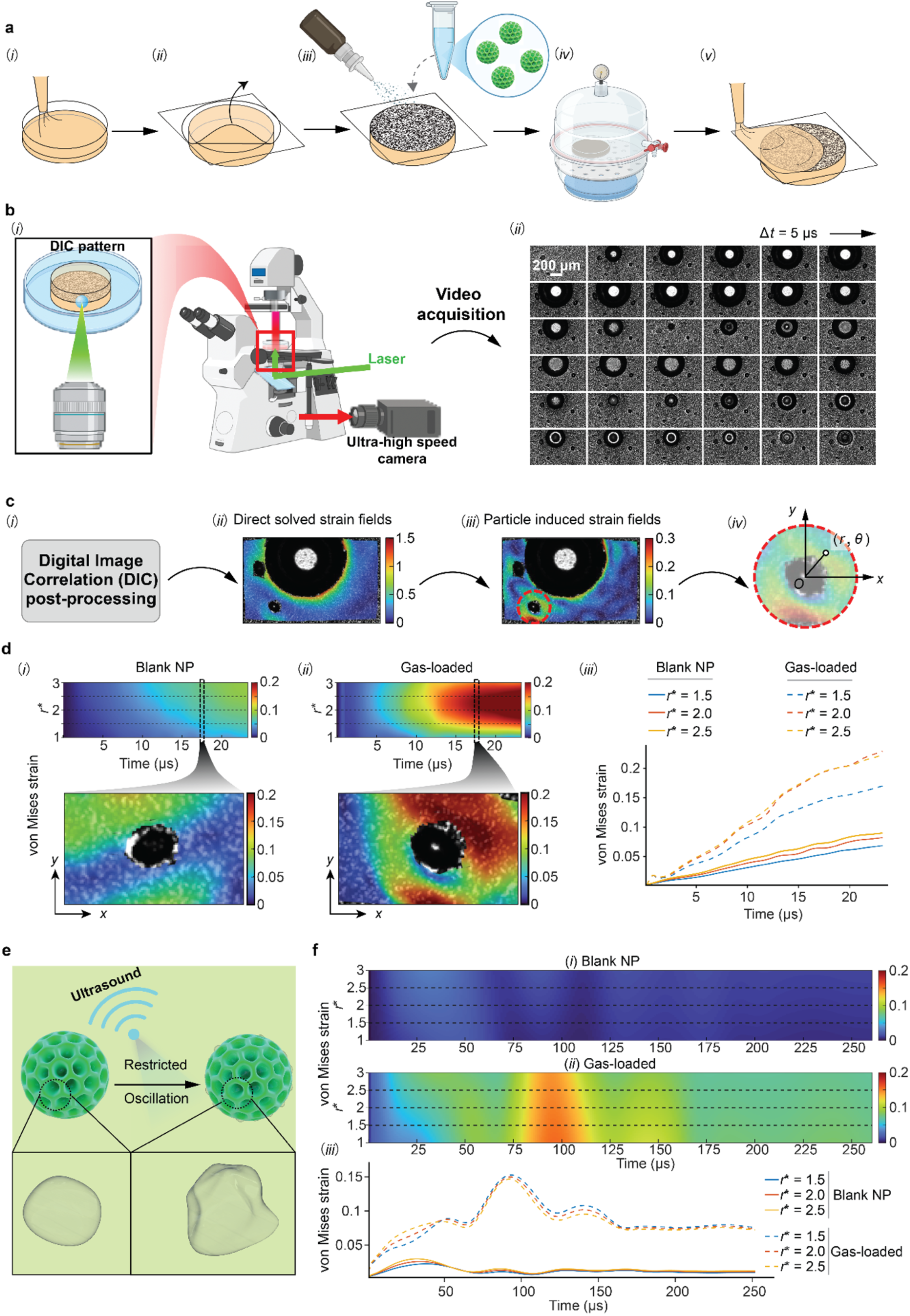
Visualization of mechanically amplified strain fields by liquid-phase gas nanotransducers. (a) Preparation of a gelatin hydrogel sample with an embedded DIC speckle pattern and liquid-phase gas nanotransducers, followed by degassing and encapsulation with a second gelatin layer. (b) (i) Experimental platform used to generate ultrasound-mimetic pressure waves via laser-induced inertial cavitation, where bubble dynamics are recorded using an ultra-high-speed camera with synchronized pulsed illumination. (ii) Selected time-resolved image sequence (Δ*t* = 5 μs) showing the nucleation, expansion, and collapse of the primary inertial cavitation bubble. (c) DIC post-processing to obtain (i) total strain fields, (ii) particle-induced strain fields obtained by subtracting the primary-bubble contribution, (iii) localized strain amplification near MOF clusters, and (iv) radial coordinate system definition used for strain quantification. (d) Spatiotemporal characterization of von Mises strain near ZIF-8 particles: (i-ii) representative von Mises strain evolution maps for blank and gas-loaded ZIF-8 particles. (iii) Radially averaged von Mises strain evolution at different normalized radial positions (*r*\* = *r*/*R*_MOF_), highlighting enhanced mechanical response in the gas-loaded particle system. (e) Schematic illustration of restricted gas oscillation within MOF pore channels under ultrasound excitation that acts as acoustic amplifiers. (f) Extended-time dynamics results acquired at 1 million frames per second (fps). (i–ii) Kymograph diagrams of particle-induced von Mises strain fields around blank and gas-loaded particles, respectively. (iii) Corresponding strain evolution at selected *r*^∗^ values, showing sustained oscillatory behavior and persistent amplification under gas-loaded conditions.

By subtracting the deformation field induced by the primary cavitation bubble, we isolate the net mechanical response associated with the nanotransducers.(*25*, *31*, *32*) As shown in **Fig. 2d, Supplementary Fig.9**, and **Supplementary Video 2**, a pronounced amplification of the local mechanical von Mises strain field is observed in the vicinity of gas-loaded MOF particles, whereas negligible responses are observed in solvent-filled controls (blank NPs). These results provide direct evidence that confined gas domains act as localized amplifiers of acoustic stress. Spatiotemporal analysis further shows that confined gas oscillations generate sustained, high-amplitude mechanical responses over multiple oscillation cycles. At the initial ultrafast timescale (0–25 μs), an intense, localized compressive deformation is rapidly induced in the surrounding matrix, consistent with confined gas–bubble oscillations. We then captured the longer-time dynamics by tracking the temporal evolution of the mechanical field over multiple oscillation cycles of the cavitation event (**Fig. 2e,f**, and **Supplementary Fig. 10**). The results reveal a sustained, pronounced excitation-induced oscillatory trend in the local strain field over an extended 256-µs observation window. Specifically, temporal strain evolution curves extracted at dimensionless radial distances *r*\* = 1, 2, and 3 (where *r*\* = r/*R*_MOF_, *r* the radius to the center of MOF, and *R*_MOF_ the MOF’s radius) confirm that the deformation amplitude driven by the gas-loaded MOFs is persistently and significantly larger than the minimal mechanical response of blank NPs. Similar behavior is observed for ZIF-67 and NU-1003 (**Supplementary Figs. 11–12**), indicating that this mechanically amplified response is a general feature of liquid-phase gas nanotransducers, akin to that of microbubbles (**Supplementary Figs. 13**).

We next quantify the acoustic output of liquid-phase gas nanotransducers using hydrophone measurements (**Fig. 3a**). As shown in **Fig. 3b**, gas-loaded ZIF-8 markedly enhances the acoustic pulse amplitude, whereas gas-free ZIF-8 shows no detectable change relative to water, establishing these liquid-phase gas nanotransducers as acoustic actuators, consistent with DIC analysis (**Fig. 2d**). Notably, the amplification index increases with ZIF-8 nanotransducer concentration and shows a trend towards equilibrium above 200 μg/mL (**Fig. 3c** and **Supplementary Fig. 14**). Similar behaviour is observed for ZIF-67 and NU-1003; however, ZIF-8 exhibits the strongest mechanical amplification effect (**Fig. 3d,e** and **Supplementary Figs. 15,16**). We speculate that this behavior may be partially attributed to its higher gas-loading capacity in aqueous media (**Fig. 1h**), which could increase the effective curvature radius of the confined gas–liquid interfaces within the pores. This, in turn, may facilitate larger-amplitude oscillations under identical acoustic excitation, thereby enhancing the observed mechanical response (detailed in **Supplementary Information**). The same concentration-dependent mechanical amplification effect was also observed in microbubbles, further supporting the role of gas–liquid interfaces in enhancing acoustic output (**Supplementary Fig. 17**).

**Fig. 3.**
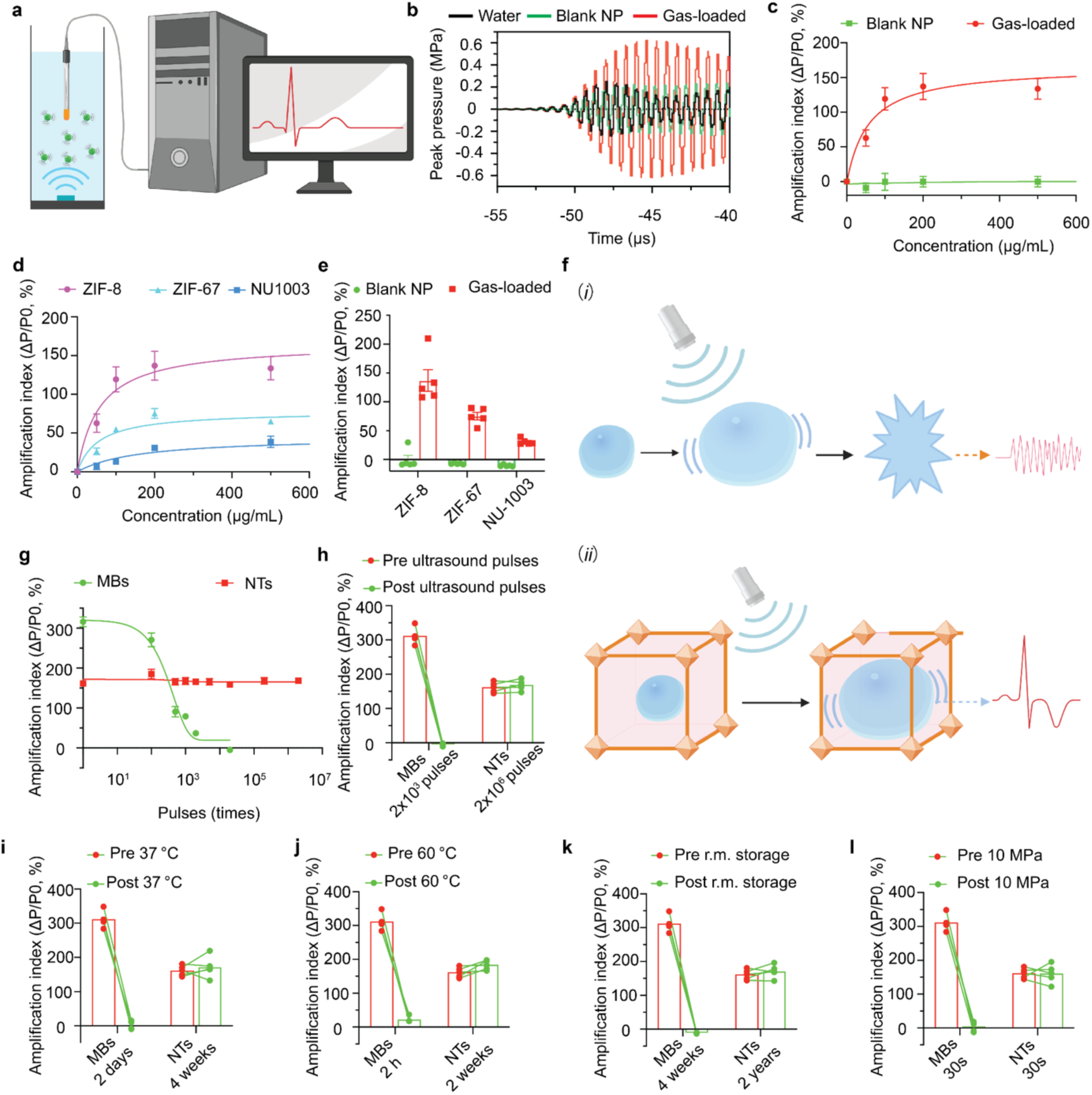
Restricted gas-bubble oscillations in liquid-phase nanotransducers enable acoustic actuation. (a) The mechanical effects of liquid-phase nanotransducers were characterized *in situ* by measuring the peak acoustic pressure using a hydrophone. (b) Acoustic waveforms recorded in aqueous solutions containing ZIF-8 nanotransducers (200 μg/mL) with or without gas loading, and in bulk water, using a 1.5 MHz focused ultrasound (FUS) transducer operated at identical output power. (c) Acoustic actuation of ZIF-8 nanotransducers with or without gas loading at varying concentrations under stimulation by a 1.5 MHz FUS transducer at identical output power. (d) Acoustic actuation of different liquid-phase gas nanotransducers, including ZIF-8, ZIF-67, and NU-1003, at varying concentrations. (e) Quantified acoustic actuation of different liquid phase gas nanotransducers at 200 μg/mL. (f) Schematic illustration showing that (i) free microbubbles oscillate and collapse under acoustic waves, resulting in loss of functionality, whereas (ii) liquid-phase gas bubbles confined within nanotransducers undergo restricted yet sustained oscillations, producing stable and amplified acoustic signals. (g) Acoustic actuation curves of ZIF-8 liquid-phase nanotransducers and commercial microbubbles under repeated acoustic stimulation. (h) Quantitative analysis comparing acoustic actuation stability of ZIF-8 liquid-phase nanotransducers and commercial microbubbles under repeated stimulation. (i-l) Quantification of acoustic actuation attenuation of microbubbles and liquid-phase nanotransducers under different conditions: (i) storage at 37 °C for 4 weeks; (j) incubation at 60 °C for 2 weeks; (k) storage at room temperature for 2 years; and (l) ultrasound treatment for 30 s at a peak pressure of 10 MPa.

Importantly, these nanotransducers operate in a fundamentally different regime from conventional gas systems.(*7*, *10*) Whereas free microbubbles undergo expansion and collapse, leading to rapid functional loss, confined gas domains within MOF pores exhibit restricted, non-destructive oscillations that sustain acoustic output (**Fig. 3f)**. As a result, liquid-phase gas nanotransducers exhibit highly stable performance over more than 2 × 10⁶ excitation cycles (300 μs on, 700 μs off; PRF, 1,000 Hz; peak pressure, 0.3 MPa), with no measurable attenuation (**Fig. 3g** and **Supplementary Fig. 18**). In contrast, conventional microbubbles fail within ∼2 × 10³ cycles due to collapse-induced gas loss (**Fig. 3h**). These results demonstrate that confined gas stabilization decouples acoustic function from bubble collapse, enabling persistent acoustic transduction.

More importantly, a major limitation of existing gas-based systems is their poor stability during storage and operation, often requiring cold-chain preservation and exhibiting rapid loss of function upon reconstitution.(*7*, *10*) In contrast, confinement of gas within hydrophobic MOF pores enables long-term stability without external stabilization strategies. ZIF-8 nanotransducers retain full acoustic functionality after 4 weeks at 37 °C, whereas microbubble solutions lose activity within 2 days (**Fig. 3i** and **Supplementary Fig. 19**). Under more stringent conditions, nanotransducers remain functional after storage at 60 °C for 2 weeks, while microbubbles fail within 2 h (**Fig. 3j** and **Supplementary Fig. 20**). Remarkably, nanotransducers maintain performance after 2 years of storage at room temperature (**Fig. 3k** and **Supplementary Fig. 21**), and withstand prolonged high-intensity ultrasound exposure without attenuation (**Fig. 3l** and **Supplementary Fig. 22**). Together, these results establish that MOF-based liquid-phase gas nanotransducers, enabled by confined gas stabilization and restricted oscillations, overcome fundamental limitations of conventional gas systems by decoupling acoustic function from bubble collapse, thereby enabling persistent acoustic transduction in liquid environments. As proof-of-concept demonstrations, we further apply this platform to ultrasound–biological interfacing, including long-term physiological monitoring and neuromodulation.

### Liquid-phase gas nanotransducers as stable acoustic reporters

Building on the non-collapsing acoustic regime established above, we next investigated liquid-phase gas nanotransducers as stable acoustic reporters for anatomical and physiological monitoring. Non-invasive visualization of the body and image-guided therapeutic intervention continues to drive advances in diagnostic imaging, where ultrasound is widely used owing to its safety, affordability, and operational flexibility.(*2*, *7*, *33*, *34*) However, conventional microbubbles are inherently unstable under acoustic excitation, as measured in phantom imaging experiments (**Supplementary Fig. 23**), where oscillations beyond a critical threshold induce collapse and rapid signal decay, thereby limiting their application in long-term ultrasound imaging (**Fig. 4a–c**). In contrast, liquid-phase gas nanotransducers operate within the non-collapsing oscillatory regime established here. Gas confined within MOF pore channels undergoes nonlinear yet stable oscillations without rupture or gas leakage, enabling sustained acoustic emission (**Fig. 3f**). This restricted oscillation is expected to preserve structural integrity and maintain consistent signal output, providing a physical basis for persistent ultrasound imaging.

**Fig. 4.**
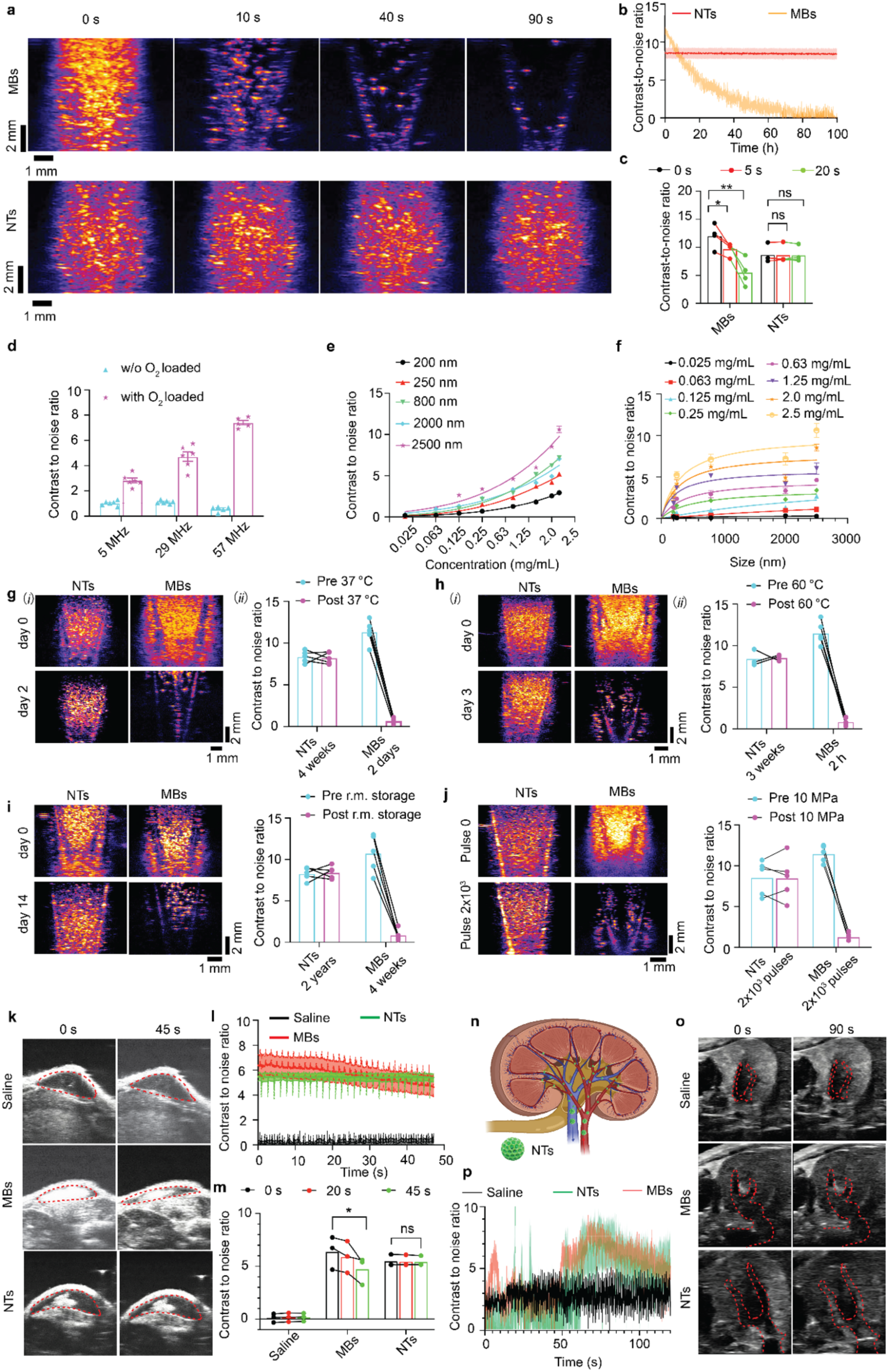
Liquid-phase gas nanotransducers function as acoustic imaging reporters. (a) Long-term ultrasound imaging of commercial microbubbles (MBs), gas vesicles (GVs), and ZIF-8 nanotransducers at 29 MHz. (b,c) Quantification of contrast-to-noise ratio (CNR) for these contrast agents and their temporal decay profiles. (d) CNR of ZIF-8 nanotransducers (800 nm) at different frequencies, including 5, 29, and 57 MHz. (e,f) CNR of ZIF-8 nanotransducers with different diameters across varying concentrations. (g) CNR changes before and after 4 weeks of incubation at 37 °C under cell-culture conditions. (h) CNR changes before and after 21 days of incubation at 60 °C. (i) CNR changes before and after 2-year storage at room temperature. (j) CNR before and after exposure to ultrasound stimulation (10 MPa). (k) *In vivo* ultrasound imaging after subcutaneous injection of MBs and ZIF-8 nanotransducers in mice. (l,m) Corresponding CNR quantification and temporal profiles. (n) Schematic of nanotransducer circulation through the kidney following intravenous administration in mice. (o) *In vivo* ultrasound imaging of renal blood flow enhanced by MBs and NTs in mice. (p) CNR quantification of renal contrast dynamics.

To validate this capability, we compared the ultrasound signal attenuation profiles of various contrast agents, including commercial microbubbles (Lumason) and oxygen-loaded ZIF-8 nanocrystals, under continuous acoustic stimulation. ZIF-8 nanotransducers were selected as a representative MOF system because they exhibited the strongest acoustic amplification and high gas-loading capacity among the tested MOFs. As shown in **Fig. 4a-c**, ZIF-8 liquid-phase gas nanotransducers maintained stable imaging signals without noticeable attenuation, whereas microbubbles exhibited rapid signal loss within approximately 40 s, attributed to their complete structural collapse. These results demonstrate that MOF liquid-phase gas nanotransducers possess exceptional signal stability under prolonged ultrasound exposure, highlighting their potential as durable, non-invasive contrast agents for real-time bioimaging applications.

We next evaluated the ultrasound imaging performance of liquid-phase nanotransducers under varying conditions using ZIF-8 as a representative system. Contrast increased with frequency, reaching a maximum at 57 MHz, where ZIF-8 liquid-phase gas nanotransducers generated ∼15-fold higher scattering than buffer controls (**Fig. 4d** and **Supplementary Fig. 24**). Moreover, ZIF-8 nanotransducers with diameters ranging from 200 to 2500 nm were synthesized, and their acoustic contrast was analyzed across different concentrations. As shown in **Fig. 4d–f** and **Supplementary Fig. 25**, the Contrast-to-Noise Ratio (CNR) increased both with particle concentration and particle size, indicating size- and dose-dependent imaging enhancement.

We next evaluated whether confined gas stabilization within MOF pores could overcome the intrinsic instability of conventional gas systems. Unlike microbubbles, which initially generate strong contrast but rapidly lose functionality owing to collapse during storage or repeated acoustic excitation, liquid-phase gas nanotransducers maintained stable acoustic performance without detectable signal attenuation under a broad range of challenging conditions (**Fig. 4g–j**). Notably, no measurable signal loss was observed after storage at 37 °C for 4 weeks, whereas microbubble signals disappeared within 2 days under the same conditions (**Fig. 4g** and **Supplementary Fig. 26**). Under even more stringent conditions, nanotransducers remained fully functional after storage at 60 °C for 2 weeks, while microbubbles again failed within 2 h (**Fig. 4h** and **Supplementary Fig. 27**). Remarkably, MOF nanotransducers preserved their acoustic functionality even after 2 years of storage at room temperature without cold-chain support (**Fig. 4i** and **Supplementary Fig. 28**), highlighting their exceptional long-term stability and potential to alleviate the logistical burden of cold-chain management for ultrasound technologies. In addition, these nanotransducers retained robust acoustic responses during prolonged high-intensity ultrasound exposure without functional degradation (**Fig. 4j**), thereby substantially extending their operational lifetime and application scope. This unusual stability is consistent with the restricted, non-destructive oscillatory dynamics of confined gas within MOF pores (**Fig. 3f–l**).

We further investigated *in vivo* performance. Following subcutaneous injection in mice, ZIF-8 nanotransducers generated robust and persistent CNR signals, remaining stable for at least 30 min (**Fig. 4k–m**). In contrast, microbubble signals progressively diminished and were nearly lost by 30 min, demonstrating the superior durability of liquid-phase nanotransducers as long-term ultrasound reporters. Finally, to assess intravenous imaging capability, we monitored renal contrast following systemic administration. Strong signals appeared in the kidney within 30 s (**Fig. 4n–p** and **Supplementary Video 3**), enabling real-time tracking of nanotransducer transit speed and direction. These results highlight the potential of liquid-phase nanotransducers for deep-tissue imaging and long-term physiological monitoring.

### Liquid-phase gas nanotransducers enable targeted ultrasound brain stimulation

Building on their ability to sustain localized acoustic amplification and stable signal output, we next investigated liquid-phase gas nanotransducers as localized acoustic actuators for long-term neuromodulation using non-invasive ultrasound. Achieving remote and spatiotemporally precise activation of neurons in targeted deep-brain regions remains a central challenge in neural circuit mapping and neurotherapeutic development. Ultrasound offers a powerful means of modulating neuronal activity in deep brain regions, particularly when coupled to mechanosensitive ion channels that confer sensitivity to mechanical stimuli.(*5*, *35–38*) However, neuronal activation in deep brain regions is limited by inefficient mechanical coupling, as a substantial fraction of acoustic energy is lost when ultrasound traverses the skull due to the pronounced impedance mismatch between bone and soft tissue, with attenuation further increasing at higher frequencies.(*39*, *40*) Skull-induced scattering further distorts the focus, displacing the focal zone and reducing targeting precision.(*39*) To overcome these limitations, we explored liquid-phase nanotransducers as ultrasound actuators capable of localizing and amplifying acoustic mechanical effects, thereby enhancing spatial precision and neuromodulation efficiency in the deep brain.

ZIF-8 liquid-phase nanotransducers were selected as a representative platform owing to their pronounced acoustic amplification properties(**Fig. 2d** and **Fig. 3c**). Their capacity to mediate ultrasound-driven neuronal modulation was evaluated in primary cortical neurons *in vitro* (**Fig. 5a**). Neurons were transduced with the genetically encoded calcium indicator GCaMP6s (AAV-9-hSyn-GCaMP6s-WPRE-SV40) and imaged in real time under varying stimulation conditions (**Fig. 5a**). Application of ultrasound in the presence of ZIF-8 nanotransducers (0.19 MPa, 1 kHz pulse repetition frequency, 50 μg/mL) induced robust neuronal responses, with ∼90% of neurons exhibiting firing at a latency of 0.70 s and multiple successive activation events (**Fig. 5b–d**, **Supplementary Fig. 29, and Supplementary Video 4**). In contrast, only sporadic activity was observed in the absence of ultrasound or with blank nanoparticles under identical ultrasound exposure (**Fig. 5c,d**). Mechanistically, neuronal activation was primarily mediated by intrinsic mechanosensitive ion channels.(*5*, *35–37*) The nanotransducers locally amplified mechanical stress, facilitating channel gating even at low ultrasound pressures, thereby markedly enhancing sonogenetic efficiency relative to ultrasound alone (**Supplementary Fig. 30**).

**Fig. 5.**
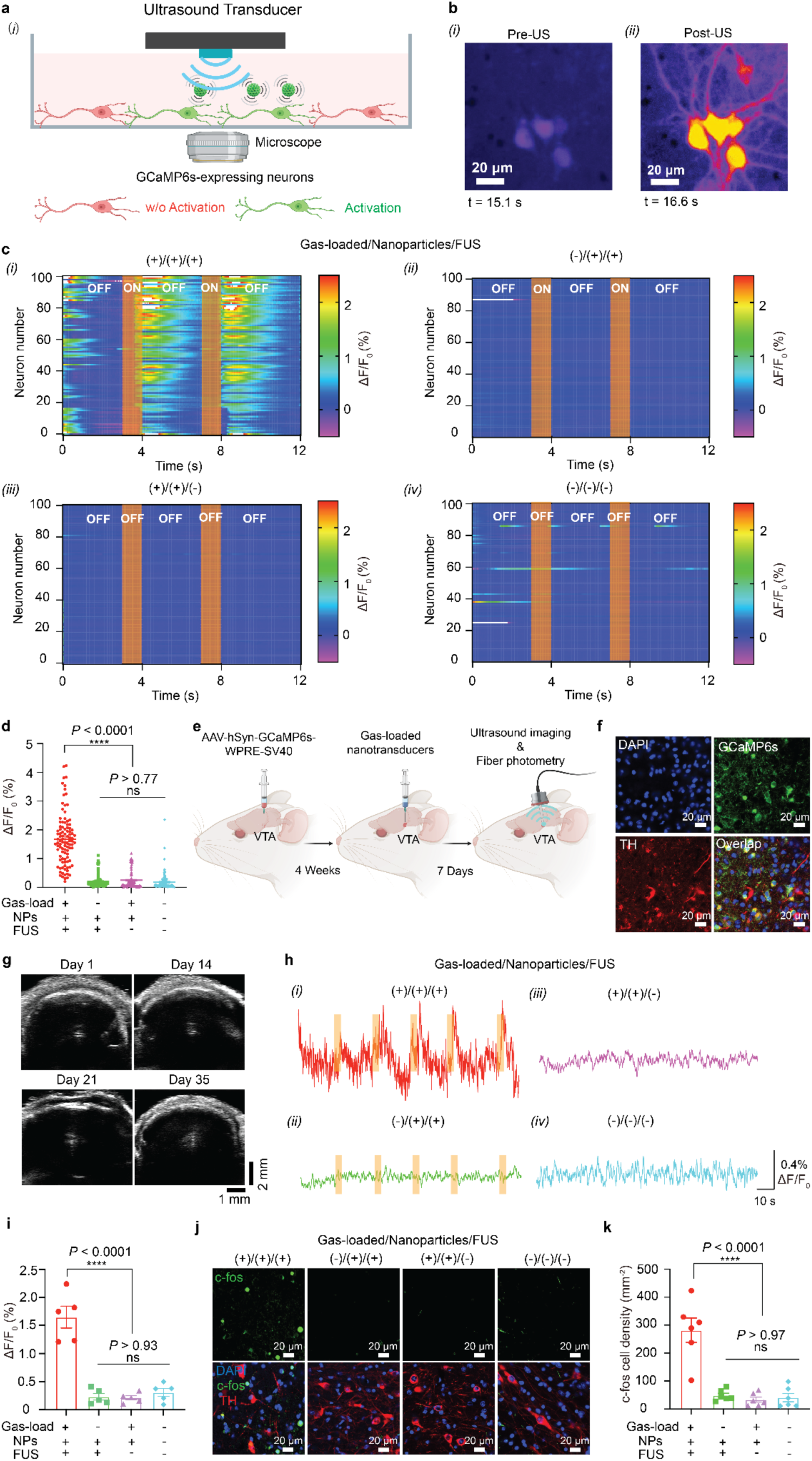
Liquid-phase gas nanotransducers function as an acoustic actuator for neuromodulation. (a) Schematic of ultrasound-triggered neuromodulation with nanotransducers, where the nanotransducers act as acoustic actuators to amplify mechanical effects under ultrasound stimulation, and then activate the neurons; (b) GCaMP6s signal in primary neurons before and after ultrasound stimulation; FUS: 1.5 MHz, 0.19 MPa, 300 ms stimulation duration, 300 ms pulse width, 40% duty cycle, 1 kHz PRF. (c) Fluorescence images of GCaMP6s-expressing neurons before and after sonogenetic stimulus. Scale bar: 20 μm (d) Heat maps of normalized GCaMP6s fluorescence intensity from 100 neurons in different experimental conditions (n = 100 neurons examined over three independent experiments for each group), including (i) gas-loaded nanotransducers with ultrasound stimulation, (ii) blank nanotransducers with ultrasound stimulation, (iii) no ultrasound stimulation, and (iv) no nanotransducers and no ultrasound stimulation. (e) Statistical analysis of calcium signal changes in 100 primary neurons under the different conditions (n = 100 neurons examined over three independent experiments for each group). Mean ± s.e.m. Two-way ANOVA and Tukey’s tests (P ≥ 0.05 (ns), ****P < 0.0001). (f) Experimental scheme of the in vivo ultrasound imaging tracking and fiber photometry in the VTA of mice during sonogenetic stimulation. (g) Confocal images showing the expression of GCaMP6s in the VTA, where the VTA area was located by TH staining of dopamine neurons. Scale bars, 20 μm. n = 3 per group. (h) Ultrasound imaging of nanotransducers after injection in VTA at various time points for sonogenetic brain stimulation. (i) Normalized GCaMP6s fluorescence change (ΔF/F_0_) in mice VTA under different sonogenetic conditions; FUS: 1.5 MHz, 0.55 MPa, 3,000 ms stimulation duration, 300 ms pulse width, 40% duty cycle, 1 kHz PRF. (j) Statistical analysis of calcium signal changes in photometry tests under different sonogenetic conditions, two-way ANOVA and Tukey’s tests (P ≥ 0.05 (ns), ****P < 0.0001). Mean ± s.e.m., n = 5 (+++), n = 5 (-++), n = 5 (++-), n = 5 (---). (k) c-fos expression in the VTA after different sonogenetic stimuli, Scale bars, 20 μm. n = 6 per group. (l) Quantification of c-fos expression in VTA after sonogenetic stimulus. Mean ± s.e.m., n = 6 per condition. Two-way ANOVA and Tukey’s tests (P ≥ 0.05 (ns), ****P < 0.0001).

We subsequently evaluated nanotransducer-enabled ultrasound neuromodulation in the ventral tegmental area (VTA) of mice using fiber photometry. The VTA, a key hub governing diverse motivational and affective behaviors, contains neurons that express endogenous mechanosensitive ion channels.(*38*, *41*) Neurons in the VTA were unilaterally transduced with AAV-9-hSyn-GCaMP6s-WPRE-SV40, followed four weeks later by stereotaxic injection of ZIF-8 liquid-phase nanotransducers into the same location (**Fig. 5e,f**). Owing to their strong acoustic responsiveness, these nanotransducers could be visualized non-invasively *in vivo*, enabling longitudinal monitoring of their presence and structural integrity. As shown in **Fig. 5g** and **Supplementary Fig. 31**, ZIF-8 nanotransducers produced pronounced ultrasound contrast at the injection site relative to blank nanoparticles and non-injected controls, and maintained robust echogenicity for over five weeks post-injection, consistent with their stability for sustained neuromodulation. We next assessed neuronal excitation induced by nanotransducer-enabled ultrasound neuromodulation by recording ultrasound-evoked Ca²⁺ influx through changes in GCaMP6s fluorescence using fiber photometry. The GCaMP6s signal increased remarkably under ultrasound stimulation with gas-loaded nanoparticles and achieved multiple times neural activation with high efficiency, but remained constant without either gas-loaded nanoparticles or ultrasound stimulation (**Fig. 5h,i**). In addition, our nanotransducer-based ultrasound neuromodulation approach demonstrated excellent stability and efficiency for long-term neural modulation even after 12 weeks of injection (**Supplementary Fig. 32**), due to their improved *in viv*o stability (**Fig. 5g**). C-fos immunofluorescence analysis, an immediate early gene and a marker for neural excitation, further demonstrated that the neurons in the VTA were targeted and only activated by ultrasound stimulus in mice with liquid-phase nanotransducer injection (**Fig. 5j,k**). We also evaluated the biocompatibility of this liquid-phase nanotransducer-based ultrasound neuromodulation approach through immunostaining brain sections. The results showed that this approach did not cause any activation of microglia (**Supplementary Fig. 33**) and astrocytes (**Supplementary Fig. 34**), nor did we observe remarkable neuron apoptosis (**Supplementary Fig. 35**), demonstrating the biosafety of this approach for remote neuromodulation.

## In conclusion

In conclusion, we establish a generalizable physical framework for stabilizing and programming gas in aqueous environments to enable persistent acoustic transduction. By confining gas within hydrophobic porous nanostructures, interfacial forces create a three-phase equilibrium that suppresses collapse while supporting reversible, non-destructive oscillations under ultrasound. This non-collapsing oscillatory regime fundamentally differs from conventional cavitation-based mechanisms and provides a physical basis for sustained acoustic energy conversion in the liquid phase. Leveraging this principle, we develop liquid-phase gas nanotransducers that function as both long-lived imaging reporters and durable acoustic actuators. Their exceptional stability—spanning prolonged storage even at elevated temperatures, and millions of excitation cycles—enables continuous ultrasound imaging and repeated neuromodulation both *in vitro* and *in vivo*, overcoming the intrinsic limitations of existing microbubble-based systems. More broadly, this work establishes gas as a programmable and persistent energy transducer in liquid environments, opening new opportunities for long-term biological interfacing, deep-tissue sensing, and non-invasive control of cellular function. These findings lay the foundation for a new class of ultrasound-driven biointerfaces and may facilitate the development of scalable, cold-chain-free technologies for biomedical imaging and therapy.

## Acknowledgment

Dr. Huiliang Wang acknowledges funding support from the National Science Foundation (NSF) CAREER award (2340964), NIH Maximizing Investigators’ Research Award (National Institute of General Medical Sciences 1R35GM147408), Robert A. Welsh Foundation Grant (No. F-2084-20210327), Craig H. Neilsen Foundation Pilot Research Grant, UT Austin Portugal Exploratory Research Projects Grant, Texas Biologics Pilot Grant and a Grant from the Alpha-1 Foundation. Dr. Noah B. Stern acknowledges support from National Institutes of Health (NIH) Training Grant (T32 EB007507). Dr. Jin Yang acknowledges funding support by the U.S. National Science Foundation (NSF) under Grants No. 2232428, 2441460, and 2452029. Dr. Jarad Mason acknowledges support from the Ono Pharma Foundation. We acknowledge BioRender.com for the figure drawing.

## Author Contributions

W.W. and H.W. designed the project. W.W. led the materials characterization, ultrasound performance studies, in vitro experiments, in vivo experiments, and data analysis. W.H. and M.W. synthesized the MOF materials and performed PXRD and BET characterizations under the supervision of J.M. Z.T., J.B., L.B., and Y.Z. conducted the DIC experiments, while Z.T. performed the DIC data analysis. J.Y. supervised and supported the design, characterization, and interpretation of the DIC studies. B.S., N.S., T.P., B.L., M.L., and P.S. assisted with ultrasound imaging and data analysis. X.L., B.A., J.H., S.Y., J.J., I.P., H.X., and M.Y. assisted with primary neuron preparation, data analysis, and figure preparation. All authors contributed to the writing and revision of the manuscript.

## Additional Information

Supplementary Information is available for this paper.

## Reference

1. J. Yan, B. Huang, J. Tonko, M. Toulemonde, J. Hansen-Shearer, Q. Tan, K. Riemer, K. Ntagiantas, R. A. Chowdhury, P. D. Lambiase, R. Senior, M.-X. Tang, Transthoracic ultrasound localization microscopy of myocardial vasculature in patients. *Nat*. Biomed. Eng. 8, 689–700 (2024).

2. A. Farhadi, G. H. Ho, D. P. Sawyer, R. W. Bourdeau, M. G. Shapiro, Ultrasound imaging of gene expression in mammalian cells. Science 365, 1469–1475 (2019).

3. E. Davoodi, J. Li, X. Ma, A. H. Najafabadi, J. Yoo, G. Lu, E. S. Sani, S. Lee, H. Montazerian, G. Kim, J. Williams, J. W. Yang, Y. Zeng, L. S. Li, Z. Jin, B. Sadri, S. S. Nia, L. V. Wang, T. K. Hsiai, P. S. Weiss, Q. Zhou, A. Khademhosseini, D. Wu, M. G. Shapiro, W. Gao, Imaging-guided deep tissue in vivo sound printing. Science 388, 616–623 (2025).

4. S. Tang, H. Han, X. Ma, P. N. Patel, C. Gong, J. Zhang, E. Criado-Hidalgo, J. Yoo, J. Li, G. Kim, S. Yin, D. Wu, M. G. Shapiro, Q. Zhou, W. Gao, Enzymatic microbubble robots. Nat. Nanotechnol., 21, 397–406 (2026).

5. X. Hou, J. Jing, Y. Jiang, X. Huang, Q. Xian, T. Lei, J. Zhu, K. F. Wong, X. Zhao, M. Su, D. Li, L. Liu, Z. Qiu, L. Sun, Nanobubble-actuated ultrasound neuromodulation for selectively shaping behavior in mice. Nat. Commun. 15, 2253 (2024).

6. A. Bar-Zion, A. Nourmahnad, D. R. Mittelstein, S. Shivaei, S. Yoo, M. T. Buss, R. C. Hurt, D. Malounda, M. H. Abedi, A. Lee-Gosselin, M. B. Swift, D. Maresca, M. G. Shapiro, Acoustically triggered mechanotherapy using genetically encoded gas vesicles. Nat. Nanotechnol. 16, 1403– 1412 (2021).

7. J. R. Lindner, Microbubbles in medical imaging: current applications and future directions. Nat. Rev. Drug Discov. 3, 527–532 (2004).

8. N. Renaudin, C. Demené, A. Dizeux, N. Ialy-Radio, S. Pezet, M. Tanter, Functional ultrasound localization microscopy reveals brain-wide neurovascular activity on a microscopic scale. Nat. Methods 19, 1004–1012 (2022).

9. Z. Jin, A. Lakshmanan, R. Zhang, T. A. Tran, C. Rabut, P. Dutka, M. Duan, R. C. Hurt, D. Malounda, Y. Yao, M. G. Shapiro, Ultrasonic reporters of calcium for deep tissue imaging of cellular signals, bioRxivorg (2023). 10.1101/2023.11.09.566364.

10. M. G. Shapiro, P. W. Goodwill, A. Neogy, M. Yin, F. S. Foster, D. V. Schaffer, S. M. Conolly, Biogenic gas nanostructures as ultrasonic molecular reporters. Nat. Nanotechnol. 9, 311–316 (2014).

11. L. G. Delogu, G. Vidili, E. Venturelli, C. Ménard-Moyon, M. A. Zoroddu, G. Pilo, P. Nicolussi, C. Ligios, D. Bedognetti, F. Sgarrella, R. Manetti, A. Bianco, Functionalized multiwalled carbon nanotubes as ultrasound contrast agents. Proc. Natl. Acad. Sci. U. S. A. 109, 16612–16617 (2012).

12. P. Frinking, T. Segers, Y. Luan, F. Tranquart, Three decades of ultrasound contrast agents: A review of the past, present and future improvements. Ultrasound Med. Biol. 46, 892–908 (2020).

13. S. Bianco, M. Hasan, A. Ahmad, S.-J. Richards, B. Dietrich, M. Wallace, Q. Tang, A. J. Smith, M. I. Gibson, D. J. Adams, Mechanical release of homogenous proteins from supramolecular gels. Nature 631, 544–548 (2024).

14. N. Giri, M. G. Del Pópolo, G. Melaugh, R. L. Greenaway, K. Rätzke, T. Koschine, L. Pison, M. F. C. Gomes, A. I. Cooper, S. L. James, Liquids with permanent porosity. Nature 527, 216–220 (2015).

15. K. Jie, Y. Zhou, H. P. Ryan, S. Dai, J. R. Nitschke, Engineering permanent porosity into liquids. Adv. Mater. 33, e2005745 (2021).

16. A. Bavykina, A. Cadiau, J. Gascon, Porous liquids based on porous cages, metal organic frameworks and metal organic polyhedra. Coord. Chem. Rev. 386, 85–95 (2019).

17. T. D. Bennett, F.-X. Coudert, S. L. James, A. I. Cooper, The changing state of porous materials. Nat. Mater. 20, 1179–1187 (2021).

18. D. P. Erdosy, M. B. Wenny, J. Cho, C. DelRe, M. V. Walter, F. Jiménez-Ángeles, B. Qiao, R. Sanchez, Y. Peng, B. D. Polizzotti, M. O. de la Cruz, J. A. Mason, Microporous water with high gas solubilities. Nature 608, 712–718 (2022).

19. D. Neumann, D. Woermann, Stability of the volume of air trapped on the abdomen of the water spider Argyroneta aquatica. Springerplus 2, 694 (2013).

20. G. Fraux, F.-X. Coudert, A. Boutin, A. H. Fuchs, Forced intrusion of water and aqueous solutions in microporous materials: from fundamental thermodynamics to energy storage devices. Chem. Soc. Rev. 46, 7421–7437 (2017).

21. V. Eroshenko, R. C. Regis, M. Soulard, J. Patarin, Energetics: a new field of applications for hydrophobic zeolites. J. Am. Chem. Soc. 123, 8129–8130 (2001).

22. F. Cailliez, M. Trzpit, M. Soulard, I. Demachy, A. Boutin, J. Patarin, A. H. Fuchs, Thermodynamics of water intrusion in nanoporous hydrophobic solids. Phys. Chem. Chem. Phys. 10, 4817–4826 (2008).

23. P. Li, S.-Y. Moon, M. A. Guelta, L. Lin, D. A. Gómez-Gualdrón, R. Q. Snurr, S. P. Harvey, J. T. Hupp, O. K. Farha, Nanosizing a metal-organic framework enzyme carrier for accelerating nerve agent hydrolysis. ACS Nano 10, 9174–9182 (2016).

24. J. Yang, V. Rubino, Z. Ma, J. Tao, Y. Yin, A. McGhee, W. Pan, C. Franck, SpatioTemporally adaptive quadtree mesh (STAQ) digital image correlation for resolving large deformations around complex geometries and discontinuities. Exp. Mech. 62, 1191–1215 (2022).

25. A. McGhee, J. Yang, E. C. Bremer, Z. Xu, H. C. Cramer III, J. B. Estrada, D. L. Henann, C. Franck, High-speed, full-field deformation measurements near inertial microcavitation bubbles inside viscoelastic hydrogels. Exp. Mech. 63, 63–78 (2023).

26. J. Noack, A. Vogel, Laser-induced plasma formation in water at nanosecond to femtosecond time scales: calculation of thresholds, absorption coefficients, and energy density. IEEE J. Quantum Electron. 35, 1156–1167 (1999).

27. X.-X. Liang, N. Linz, S. Freidank, G. Paltauf, A. Vogel, Comprehensive analysis of spherical bubble oscillations and shock wave emission in laser-induced cavitation. J. Fluid Mech. 940 (2022).

28. D. Ranjan, J. Oakley, R. Bonazza, Shock-bubble interactions. Annu. Rev. Fluid Mech. 43, 117– 140 (2011).

29. M. Adama Maiga, O. Coutier-Delgosha, D. Buisine, A new cavitation model based on bubble-bubble interactions. Phys. Fluids (1994) 30, 123301 (2018).

30. N. A. Pelekasis, A. Gaki, A. Doinikov, J. A. Tsamopoulos, Secondary Bjerknes forces between two bubbles and the phenomenon of acoustic streamers. J. Fluid Mech. 500, 313–347 (2004).

31. J. B. Estrada, C. Barajas, D. L. Henann, E. Johnsen, C. Franck, High strain-rate soft material characterization via inertial cavitation. J. Mech. Phys. Solids 112, 291–317 (2018).

32. J. Yang, H. C. Cramer III, C. Franck, Extracting non-linear viscoelastic material properties from violently-collapsing cavitation bubbles. Extreme Mech. Lett. 39, 100839 (2020).

33. H. Tang, Y. Yang, Z. Liu, W. Li, Y. Zhang, Y. Huang, T. Kang, Y. Yu, N. Li, Y. Tian, X. Liu, Y. Cheng, Z. Yin, X. Jiang, X. Chen, J. Zang, Injectable ultrasonic sensor for wireless monitoring of intracranial signals. Nature 630, 84–90 (2024).

34. J. Liu, N. Liu, Y. Xu, M. Wu, H. Zhang, Y. Wang, Y. Yan, A. Hill, R. Song, Z. Xu, M. Park, Y. Wu, J. L. Ciatti, J. Gu, H. Luan, Y. Zhang, T. Yang, H.-Y. Ahn, S. Li, W. Z. Ray, C. K. Franz, M. R. MacEwan, Y. Huang, C. W. Hammill, H. Wang, J. A. Rogers, Bioresorbable shape-adaptive structures for ultrasonic monitoring of deep-tissue homeostasis. Science 383, 1096– 1103 (2024).

35. S. Cadoni, C. Demené, I. Alcala, M. Provansal, D. Nguyen, D. Nelidova, G. Labernède, J. Lubetzki, R. Goulet, E. Burban, J. Dégardin, M. Simonutti, G. Gauvain, F. Arcizet, O. Marre, D. Dalkara, B. Roska, J. A. Sahel, M. Tanter, S. Picaud, Ectopic expression of a mechanosensitive channel confers spatiotemporal resolution to ultrasound stimulations of neurons for visual restoration. Nat. Nanotechnol., 18, 667–676 (2023).

36. Y. Yang, J. Yuan, R. L. Field, D. Ye, Z. Hu, K. Xu, L. Xu, Y. Gong, Y. Yue, A. V. Kravitz, M. R. Bruchas, J. Cui, J. R. Brestoff, H. Chen, Induction of a torpor-like hypothermic and hypometabolic state in rodents by ultrasound. Nat. Metab. 5, 789–803 (2023).

37. M. Duque, C. A. Lee-Kubli, Y. Tufail, U. Magaram, J. Patel, A. Chakraborty, J. Mendoza Lopez, E. Edsinger, A. Vasan, R. Shiao, C. Weiss, J. Friend, S. H. Chalasani, Sonogenetic control of mammalian cells using exogenous Transient Receptor Potential A1 channels. Nat. Commun. 13, 600 (2022).

38. Y. Matsushita, K. Yoshida, M. Yoshiya, T. Shimizu, S. Tsukamoto, N. Kudo, Y. Takeuchi, M. Higuchi, M. Shimojo, TRPC6 is a mechanosensitive channel essential for ultrasound neuromodulation in the mammalian brain. Proc. Natl. Acad. Sci. U. S. A. 121, e2404877121 (2024).

39. T. S. Riis, T. D. Webb, J. Kubanek, Acoustic properties across the human skull. Ultrasonics 119, 106591 (2022).

40. W. Wang, Y. Shi, W. Chai, K. W. K. Tang, I. Pyatnitskiy, Y. Xie, X. Liu, W. He, J. Jeong, J.-C. Hsieh, A. R. Lozano, B. Artman, X. Shi, N. Hoefer, B. Shrestha, N. B. Stern, W. Zhou, D. W. McComb, T. Porter, G. Henkelman, B. Chen, H. Wang, H-bonded organic frameworks as ultrasound-programmable delivery platform. Nature 638, 401–410 (2025).

41. J. Wang, M. Su, D. Zhang, L. Zhang, C. Niu, C. Li, S. You, Y. Sang, Y. Zhang, X. Du, H. Zhang, The cation channel mechanisms of subthreshold inward depolarizing currents in the mice VTA dopaminergic neurons and their roles in the chronic-stress-induced depression-like behavior. Elife 12 (2024).

