## Supplementary Materials for "Stabilizing gas in aqueous liquids with metal–organic frameworks for persistent acoustic transduction"

**Synthesis of ZIF-8 nanocrystals**.

ZIF-8 nanocrystals were synthesized following previously reported methods with minor modifications. (*1*) Briefly, zinc nitrate hexahydrate [$Zn(NO_3)_2·6H_2O$, 1.0 g, 3.4 mmol] and 2-methylimidazole (2.24 g, 27.3 mmol) were each dissolved in 30 mL of methanol. The 2-methylimidazole solution was then added to the zinc nitrate solution under vigorous stirring (1000 rpm) at room temperature. After 1 hour of stirring, the initially clear solution became turbid, indicating the formation of ZIF-8 nanocrystals. The resulting nanocrystals were collected by centrifugation at 12,000 rpm (13,523 × g) for 10 minutes. The precipitate was washed three times with methanol to remove unreacted precursors and dried under vacuum. The dried ZIF-8 nanocrystal powder was characterized by X-ray diffraction (XRD) and subsequently redispersed in methanol at a concentration of 10 mg/mL for further experiments.

**Synthesis of ZIF-67 nanocrystals**.

ZIF-67 nanocrystals were synthesized using a similar procedure to that of ZIF-8, with cobalt nitrate replacing zinc nitrate. (*1*) Specifically, cobalt(II) nitrate hexahydrate [$Co(NO_3)_2·6H_2O$, 1.0 g, 3.4 mmol] and 2-methylimidazole (2.24 g, 27.3 mmol) were each dissolved in 30 mL of methanol. The 2-methylimidazole solution was added to the cobalt nitrate solution under vigorous stirring (1000 rpm) at room temperature. After 1 hour, the formation of ZIF-67 nanocrystals was indicated by a turbid blue coloration. The nanocrystals were collected by centrifugation at 12,000 rpm (13,523 × g) for 10 minutes, washed three times with methanol, and dried under vacuum. The dried ZIF-67 powder was characterized by XRD and redispersed in methanol at 10 mg/mL for subsequent experiments.

**Synthesis of MOF-808 nanocrystals.**

Trimesic acid (210 mg, 1.0 mmol) and $ZrOCl_2·8H_2O$ (970 mg, 3.0 mmol) were dissolved in a mixed solvent of formic acid (45 mL) and N,N-Dimethylformamide (DMF, 45 mL) in a screw-cap glass bottle. The reaction mixture was heated at 130 °C for 48 hours in an oven. After cooling to room temperature, the resulting white solid was collected by centrifugation at 12,000 rpm (13,523 × g) and sequentially washed with DMF, distilled water, and acetone (50 mL × 3 for each solvent). The solid was dried in an oven at 60 °C overnight. The dried product (800 mg) was then immersed in 1 M HCl solution (100 mL) at room temperature for 24 hours to exchange formate ligands. The solid was collected by centrifugation, washed with distilled water and acetone (50 mL × 3 each), and finally dried at 60 °C overnight to yield MOF-808 as a white powder (750 mg). The dried MOF-808 powder was characterized by XRD and redispersed in methanol at 10 mg/mL for subsequent experiments.

**Synthesis of NU-1000 nanocrystals.**

NU-1000 was synthesized following a slightly modified procedure. Briefly, $ZrOCl_2 \cdot 8H_2O$ (97 mg, 0.30 mmol) and benzoic acid (2700 mg, 22 mmol) were combined in 8 mL of DMF (in a 6-dram vial) and ultrasonically dissolved. The clear solution was heated in an oven at 80°C for 1 hour. After cooling to room temperature, $H_4TBAPy$ (40 mg, 0.06 mmol) was added to the solution, and the mixture was sonicated for 20 minutes. The yellow suspension was then heated in an oven at 80°C for 24 hours. The resulting yellow polycrystalline product was isolated by filtration and washed with DMF (10 mL × 3) over 24 hours. The solid was then suspended in 12 mL of DMF and 0.5 mL of 8 M HCl, stirred at room temperature for 24 hours, and washed with DMF (10 mL × 3) and acetone (10 mL × 3). Centrifugation (7000 rpm for 5 minutes) was used to separate the liquid and solid phases between washings. The final product was dried at room temperature under vacuum until a constant weight was achieved. The dried NU-1000 powder was characterized by XRD and redispersed in methanol at 10 mg/mL for subsequent experiments.

**Synthesis of NU-1003 nanocrystals.**

Solution A was prepared by dissolving $ZrOCl_2 \cdot 8H_2O$ (160 mg) and benzoic acid (2.0 g) in 40 mL of N,N-dimethylformamide (DMF) within a 250 mL glass bottle. The solution was heated to 80 °C for 1 hour under continuous stirring and then allowed to cool to room temperature. Subsequently, 64 mg of the organic linker 1,3,6,8-tetrakis(4-carboxy-naphthalen-2-yl)pyrene was dissolved in an additional 40 mL of DMF and added to the solution, ensuring complete dissolution. The resulting stock solution A was treated with 1,500 μL of trifluoroacetic acid (44.16 mmol) and subjected to sonication for 10 minutes. The reaction mixture was then placed in an oven at 120 °C for 3 hours. The resulting light-yellow NU-1003 product was collected by centrifugation, washed thoroughly with DMF (3 × 20 mL) and acetone (3 × 20 mL), and subsequently soaked in fresh acetone (20 mL) for 6 hours. For the activation of NU-1003, the obtained solid was soaked in a mixture of 100 mL DMF and 10 mL of 8 M HCl solution at 100 °C for 10 hours. Following this, the MOF solid was extensively washed with DMF (3 × 20 mL) and ethanol (3 × 20 mL). Finally, the solid was dried at 120 °C under vacuum for 8 hours. The dried NU-1003 powder was characterized by XRD and redispersed in methanol at 10 mg/mL for subsequent experiments.

**Gas-loaded MOFs nanoparticle solution preparation**.

A total of 1 mL of MOF methanol solution (10 mg/mL) was added to a 100 mL round-bottom flask, and the methanol was removed by vacuum distillation at 50 °C for 1 h to form a thin film. The flask was then filled with a gas ($CO_2$, $O_2$, $N_2$, or air) and evacuated; this fill–pump cycle was repeated three times. Subsequently, 2 mL of distilled water was added, and the film was sonicated for 2 min to resuspend the MOFs. Next, 2 mL of Pluronic F127 solution (20 mg/mL) was added, and the mixture was stirred at 500 rpm for 4 h at room temperature. The resulting nanoparticles were collected by centrifugation at 12,000 rpm (13,523 × g, 5 min), washed three times with distilled water, and stored in aqueous suspension at 10 mg/mL. The F127 coating improves colloidal stability and prevents nanoparticle aggregation in solution. The final suspension was used for dynamic light scattering (DLS) measurements at 0.5 mg/mL and for transmission electron microscopy (TEM) sample preparation.

For gas-free MOF nanoparticles (blank nanoparticle) preparation, 1 mL of MOF methanol solution (10 mg/mL) was added to 5 mL of Pluronic F127 solution (20 mg/mL) under stirring at 500 rpm for 4 h at room temperature. The resulting nanoparticles were then collected by centrifugation at 12,000 rpm (13,523 × g, 5 min), washed three times with distilled water, and finally resuspended and stored in aqueous solution at 10 mg/mL.

**Solid-state gas adsorption measurements of MOFs.**

Solid powders were loaded into glass sample tubes and activated under vacuum using a Micromeritics SmartVacPrep equipped with a turbomolecular pump. MOF samples were activated at room temperature for NU-1003 and at 120 °C for ZIF-8, ZIF-67, and NU-1000. MOF-808 was activated at 40 °C for 2 h, 80 °C for 6 h, and 100 °C overnight. Adsorption data were then recorded using a Micromeritics MicroActive 3Flex 3500. BET surface areas for each material were obtained from $N_2$ adsorption isotherms at 77 K. Isotherm measurements at 20 °C and 37 °C were performed using a recirculating dewar connected to an isothermal bath.

**Oxygen-release measurements in deoxygenated water of MOFs nanoparticles.**

Water was added directly to pre-massed vials of solid powders and sonicated to disperse MOF solutions. Oxygen release into deoxygenated water was quantified with a Unisense MicroRespiration $O_2$ Microsensor. Specifically, nanopure water was deoxygenated by sparging with $N_2$ for at least 30 min. A two-point calibration was performed by placing the sensor in fully nitrogenated water to obtain the mV reading corresponding to a concentration of 0 mg/L and then placing the sensor in air-equilibrated water of a known temperature to calibrate the corresponding mV reading to the known mg/L (obtained from an $O_2$ concentration table). After calibration, an aliquot of deoxygenated water was then sealed in a 1.2 mL air-tight glass chamber with a gas-tight port and a glass stir bar. The chamber was submerged in a room temperature water bath containing a temperature probe, and the sensor was inserted into the top port to record a baseline $O_2$ concentration (mg/L). Then, a known volume of an air-equilibrated MOF solution was injected into the measurement chamber through the port, and the resultant change in mg/L was measured to calculate the amount of $O_2$ released into water. The chamber was submerged in a room temperature water bath containing a temperature probe, and an automatic temperature correction of the measured $O_2$ concentrations was applied by the $O_2$ sensor software. Calculations of oxygen-release data were subsequently performed following a previous report.(*1*)

**Contact Angle Measurements of MOFs.**

MOF tablets were prepared using a tablet compression machine (Regular Pressure Gauge PR0316A) at a pressure of 7.5 tons. The resulting tablets were subjected to contact angle measurements using an FTA200 contact angle goniometer. Briefly, a 5 μL droplet of distilled water was placed onto the tablet surface, and the droplet profile was recorded by the integrated camera. The contact angle was subsequently calculated from the captured images.

**Laplace analysis of confined gas stability within MOF pore channels**

The ability of microporous MOFs to capture and stabilize gas within hydrophobic pore channels has been demonstrated to be a thermodynamic process.(*1*) In aqueous environments, randomly arranged water molecules in the bulk phase cannot spontaneously penetrate confined hydrophobic pore channels, as this would require adopting a more ordered and less mobile configuration, which is entropically unfavourable and therefore non-spontaneous.(*1*) Experimental studies have shown that pressures of at least 200 bar are needed to force water molecules into the micropores of hydrophobic zeolitic imidazolate frameworks.(*2*) Although gas stabilization within hydrophobic microporous MOFs has been experimentally validated, a general theoretical framework capable of guiding the rational design of microporous materials—not limited to MOFs—as liquid-phase gas nanotransducers for biomedical applications remains lacking. In particular, we sought to establish a model describing the three-phase equilibrium of confined gas within porous structures, thereby guiding the design of porous materials for liquid-phase gas delivery and programmable acoustic transduction under ultrasound stimulation. To this end, the law of Laplace was applied to describe the physical stability of confined gas within porous channels.

The law of Laplace is expressed as follows (see **Fig. 1c** in main text):

$$\Delta P = -\frac{2\gamma \cos\theta}{r} \tag{S1}$$

$$\Delta P = P_{\mathrm{w}} - P_{\mathrm{a}} \tag{S2}$$

$$R_{\mathrm{e}} = \frac{r}{|\cos\theta|} \tag{S3}$$

$$\theta = 180^{\circ} - \alpha \tag{S4}$$

$$P_{\mathrm{w}} = \rho_{\mathrm{w}} g h \tag{S5}$$

Here, $P_{\mathrm{w}}$ denotes the hydrostatic pressure of the surrounding water at the position of the MOFs, whereas $P_{\mathrm{a}}$ represents the pressure of the gas confined within the MOF pore channels. $\gamma$ is the air–water interfacial tension, and $r$ is the radius of the gas bubbles captured in the pore channel. $R_{\mathrm{e}}$ denotes the radius of curvature of the air–water interface spanning the two three-phase contact points. $\theta$ represents the contact angle of the MOF surface, $\rho_{\mathrm{w}}$ is the density of water, $g$ is the gravitational acceleration constant, and $h$ denotes the immersion depth of the MOFs in water.

We consider MOFs containing gas confined within pore channels in aqueous solution. In the equilibrium state (denoted by subscript e), the MOFs are positioned below the air–water interface and spontaneously return to this state after perturbation by external forces, such as ultrasound-induced stress or elevated hydrostatic pressure. The physical stability of gas confined within MOF pore channels can be understood by analyzing the balance of interfacial forces acting on the trapped gas domains. Specifically, the confined gas experiences three competing forces: inward hydrostatic pressure ($P_w$), outward gas pressure ($P_a$), and interfacial tension (**Fig. 1d**), the direction of which is determined by the surface properties of the MOF inner walls. For hydrophobic pore surfaces ($\theta > 90°$, $\alpha < 90°$, $\cos\theta < 0$), interfacial tension acts outward, opposing water intrusion into the micropores and thereby stabilizing gas within the pore channels. Under these conditions, water can displace the confined gas only when sufficiently high external pressures disrupt the interfacial equilibrium; for example, pressures exceeding ~200 bar are required to force water into the micropores of zeolitic imidazolate frameworks.(*2*, *3*) By contrast, for hydrophilic pore surfaces ($\theta < 90°$, $\alpha > 90°$, $\cos\theta > 0$), interfacial tension acts inward to promote spontaneous water penetration into the pore channels, thereby destabilizing confined gas. Based on this model, we can explain why hydrophobic porous MOFs preferentially remain dry and resist water intrusion.

Next, we further developed this model to explain how gas-loading behaviour influences the acoustic performance of liquid-phase gas nanotransducers in aqueous environments. According to Equation 1, $\Delta P$ represents the pressure redundancy resisting perturbation by external forces, where larger $\Delta P$ values correspond to stronger resistance against water intrusion into the pore channels. As $\Delta P$ decreases, the confined gas domains become increasingly susceptible to deformation under external stimulation. Increasing the gas-loading capacity effectively enlarges the confined gas volume and interfacial curvature radius ($r$), thereby facilitating gas deformation and oscillation under ultrasound stimulation. Larger interfacial curvature radii further yield enhanced mechanical amplification during oscillation, which may explain why ZIF-8 exhibits the strongest acoustic mechanical amplification among the investigated porous materials (**Fig. 3d,e**).

Following Equation 1, $\gamma_{a/w}$ is the air–water interfacial tension ($\gamma_{a/w} = 72.5 \times 10^{-7}$ J cm$^{-2}$ at 25 °C)(*4*), the pore cavity diameter of ZIF-8 is ~1.16 nm,(*1*) and the average particle size is ~800 nm. When gas is confined throughout these porous particles, the effective confined gas domain size ($2r$) spans from the nanoscale pore dimension (~1.16 nm) to the microscale particle dimension (~400 nm radius). Given the contact angle of ZIF-8 ($\theta = 94.8°$), the calculated $\Delta P$ ranges from ~0.31 to 242.99 J·cm$^{-3}$, corresponding to pressures of approximately 3.1~2429.9 bar required to force water into the confined pore structures. These values are consistent with previous experimental observations showing that pressures exceeding ~200 bar are required to force water into the micropores of hydrophobic zeolitic imidazolate frameworks.(*2*, *3*)

These results explain why liquid-phase gas nanotransducers remain stable in aqueous solution while sustaining restricted oscillations under ultrasound stimulation(**Fig. 1h** and **Fig. 3f**), why nanotransducers with higher gas-loading capacities exhibit greater acoustic amplification (**Fig. 3d,e**), and why acoustic contrast increases with particle size during ultrasound imaging (**Fig. 4e**). More broadly, this framework establishes design principles for tailoring porous materials through pore geometry and surface wettability to programmably regulate their acoustic performance.

***In vitro* ultrasound imaging.**

*In vitro* ultrasound imaging tests were conducted using a custom-fabricated silicone phantom. As illustrated in **Supplementary Fig.22**, a mold was first designed to create the phantom structure. Ecoflex™ 00-30 silicone rubber (Smooth-On, Inc.) was prepared by mixing Part A and Part B in a 1:1 weight ratio and then poured into the mold. The mixture was left to cure overnight at room temperature to form the final phantom. The resulting phantom consisted of an array of 6 × 6 conical wells, each with a diameter of 8 mm and a depth of 15 mm. For imaging tests, approximately 0.3 mL of the test solution was added to each well before ultrasound scanning with the Vevo F2 LAZR-X (FUJIFILM VisualSonics) at 5 MHz, 29 MHz, and 57 MHz central frequency. The ultrasound transducer was placed over the phantom and contacted with the solution drop for scanning. B-mode images were acquired for ultrasound imaging performance evaluation.

**Imaging data analysis**.

The ultrasound imaging performance of liquid-phase gas nanotransducers was evaluated using the contrast-to-noise ratio (CNR), following previously reported methods. Briefly, acquired ultrasound images were first analyzed using ImageJ software (ImageJ; Process → Math → Log). Rectangular regions of interest (ROIs) were then manually selected over the lesion area and the background, respectively. The mean intensity ⟨L⟩ and variance ($\sigma^2$) of the log-transformed images within each ROI were calculated. The contrast-to-noise ratio (CNR) was determined using the following equation:

$$CNR = \frac{\langle L_{\mathrm{lesion}} \rangle - \langle L_{\mathrm{background}} \rangle}{\sqrt{\sigma^2_{\mathrm{lesion}} + \sigma^2_{\mathrm{background}}}} \tag{S6}$$

where $\langle L_{(\cdot)} \rangle$ and $\sigma^2_{(\cdot)}$ are the mean and variance of the log image.

**Acoustic Amplification Evaluation of Gas-Loaded MOF Solutions.**

The mechanical amplification effects of gas-loaded MOFs were assessed through in situ acoustic peak-pressure measurements (**Fig. 3a**). Briefly, an ultrasound transducer was fixed at the bottom of a tube, and a hydrophone (HGL-0200, Onda Corporation) was positioned at the acoustic focus. The transducer was driven by an ultrasonic energy emitter (Image Guided Therapy System) to generate acoustic waves. The hydrophone was connected to an oscilloscope (SDS1204-XE, Siglent) through a signal preamplifier (AG-2010, Onda), and the output waveforms were recorded and processed using a custom MATLAB script. Gas-loaded or unloaded MOF nanoparticle suspensions were added to the tube to immerse both the transducer and hydrophone fully. Acoustic peak pressure was quantified using a single ultrasound pulse (300 µs on, 700 µs off) at a center frequency of 1.5 MHz.

**Digital Image Correlation (DIC) random speckle pattern preparation.**

To prepare a 2D planar Digital Image Correlation (DIC) speckle pattern, we employed the embedded speckle pattern method following the protocol developed by McGhee et al. (*5*, *6*) We first synthesized a flat, thin layer of gelatin hydrogel with a mass concentration of 10 wt% using porcine gelatin (300-g bloom, Sigma-Aldrich G1890) mixed with Milli-Q deionized (DI) water. The gelatin-water mixture was heated above 60 ∘C and stirred continuously until all gelatin was dissolved. The gelatin solution was then pipetted into the bottom of a 14 mm glass-bottom petri dish (VWR 10810-054) to form a layer approximately 1 mm thick. A polyethylene terephthalate (PET) plastic film was gently placed on top to create a flat, horizontal surface. The sample was then placed in an ice bath for 30 minutes to allow complete gelation. After gelatin, the PET film was carefully peeled from one edge to avoid damaging the prepared flat gelatin surface.

Micrometer-sized inkjet toner particles were then deposited onto the gelatin surface using a high-pressure air stream to minimize exposure time to prevent dehydration of the gel surface. Subsequently, the metal-organic framework (MOF) particle suspension was diluted to 100 µg/mL, and an ultrathin layer of the diluted suspension was pipetted onto the speckle-patterned surface.

Due to interparticle interactions, the MOF particles attracted nearby toner particles, forming localized microscale clusters suitable for DIC tracking. In our laser-induced cavitation experiments, we typically observed 3-20 MOF particles within the field of view (FOV), which measured 400 pixels × 250 pixels, corresponding to a physical size of 1.28 mm × 0.8 mm.

To remove air bubbles unintentionally introduced during deposition of the MOF suspension, the sample was placed in a vacuum desiccator at −25 mmHg for 30 min. Based on our experimental observations, this degassing step did not cause gas release from the MOF particles. The petri dishes were loosely covered during degassing to reduce dehydration. Immediately following degassing, an additional layer of 10 wt% gelatin solution, preheated to approximately 35 °C above its melting temperature to ensure complete liquefaction, was gently poured over the MOF-particle layer to encapsulate the speckle pattern. The thickness of this second gelatin layer exceeded 2 mm.

Using this procedure, we prepared six sample types corresponding to three different MOF materials (ZIF-8, ZIF-67, and NU-1003), each under two conditions: gas-loaded and unloaded. As a comparison case, commercial microbubbles (Lumason®, 1 × $10^3$ microbubbles/mL; kindly provided by Dr. Pengfei Song's laboratory) were also embedded within the speckle-pattern layer in gelatin using the same encapsulation protocol.

**Laser-induced inertial microcavitation**

We employed our previously developed laser-induced inertial microcavitation (LIC) experimental platform that was described in detail in prior publications.(*7*, *8*) Briefly, a tunable (1-25 mJ) Q-switched Nd:YAG Minilite II laser (Continuum, Milpitas, CA) with a pulse duration of 3–5 ns was frequency-doubled to 532 nm, expanded, and directed into the rear aperture of a Nikon Plan Fluor 10×/0.3 NA infinity-corrected objective through the backport of a Nikon Ti2-E inverted microscope (Nikon Instruments, Long Island, NY). The pulsed laser was focused onto the DIC speckle-pattern plane, corresponding to the same depth at which the MOF particles were embedded.

Absorption of the laser pulse at the focal point induce optical breakdown and nucleate a primary inertial cavitation bubble that underwent rapid expansion followed by violent collapse.(*9*, *10*) This process generate a strong shock wave that propagated radially outward from the laser focus into the surrounding hydrogel.(*11*, *12*) The primary bubble exhibits transient radial oscillations characterized by expansion–collapse cycles, producing highly localized, high–strain-rate deformation fields.(*13*) The emitted shock wave and the subsequent oscillatory motion of the primary bubble dynamically interacted with nearby MOF clusters, inducing secondary radial oscillations through fluid–structure coupling and bubble–particle interaction mechanisms.(*14–16*) These interactions are governed by local pressure gradients, added-mass effects, and transient stress-wave loading.

The temporal evolution of both the primary cavitation bubble and the MOF clusters was recorded using an ultra-high-speed camera (HPV-X2, Shimadzu Corporation, Kyoto, Japan) operating at 1–10 million frames per second (fps). Frame-by-frame illumination was provided by a synchronized SILUX640 laser illumination system (Shimadzu Corporation, Kyoto, Japan) with a pulse duration of 10-20 ns, coupled through the microscope optical path and triggered by the HPV-X2 to ensure precise temporal synchronization.

Distinct dynamic responses were observed among different MOF clusters and comparison microbubbles, reflecting variations in their effective mechanical properties, including cluster size, stiffness, viscoelasticity, and internal gas content. These differences directly influenced their oscillatory amplitude, phase response, and strain amplification under the transient shock loading generated by laser-induced cavitation.

**Mechanical stress field analysis.**

By comparing the image frames during cavitation with the reference frame and applying our recently developed SpatioTemporally Adaptive Quadtree mesh Digital Image Correlation (STAQ-DIC) image tracking algorithm,(*17*, *18*) we successfully reconstruct the history of full-field material deformations within a planar slice during each LIC event.(*19*) In STAQ-DIC, we employ an incremental tracking mode where every two consecutive frames are compared to each other to obtain an incremental displacement field. All of the images are 400 pixels × 250 pixels in dimension. We utilized the adaptive quadtree mesh in our DIC post-processing. The DIC mesh was refined near the laser-induced primary bubble and MOF clusters to enhance the DIC measurement spatial resolution. The coarsest DIC element size (DIC window spacing) in the generated adaptive quadtree mesh here is 8 pixels × 8 pixels, and the finest mesh element size is 2 pixels × 2 pixels. All the tracked displacement components are further used to calculate the full-field strain components by using a virtual strain gauge size of 40 pixels × 40 pixels. For the area surrounding each MOF cluster, its deformation is the summation of a deformation due to the primary bubble and its own reactivated dynamics. For each MOF cluster's local area, we subtract the dynamics due to the primary bubble and isolate the net mechanical response associated with the MOF nanotransducers.

**Physics of laser-induced cavitation and emitted pressure waves to excite MOF particles**

To elucidate the strain amplification observed around liquid-phase gas nanotransducers, the underlying physics can be interpreted as two coupled dynamical stages. First, a single nanosecond laser pulse induces a primary cavitation bubble that rapidly nucleates, expands, and undergoes violent collapse. The temporal evolution of the bubble radius $R(t)$ can be described by the Gilmore equation,(*10*, *20*) which accounts for liquid compressibility.

$$\left(1-\frac{\dot{R}}{c}\right)R\ddot{R}+\frac{3}{2}\left(1-\frac{\dot{R}}{3c}\right)\dot{R}^2=\left(1+\frac{\dot{R}}{c}\right)(H_\mathrm{b}-H_\infty)+\frac{R}{c}\frac{\mathrm{d}}{\mathrm{d}t}(H_\mathrm{b}-H_\infty), \tag{S7}$$

where $c$ is the local sound speed at the bubble wall, and $H$ denotes the enthalpy, defined as

$$H=\int_{p_\infty}^{p_\mathrm{b}}\frac{\mathrm{d}p}{\rho}. \tag{S8}$$

Subscripts “b” and “∞” mean the values at the bubble wall and the infinite far field, respectively. The compressible dynamics captured by the Gilmore model enable accurate description of the high-pressure impulses and shock wave emission generated during bubble nucleation and collapse. These high-amplitude, short-duration pressure pulses act as strong transient driving forces, consistent with the intense localized compressive strains observed experimentally within the first 0–25 μs.Following the initial collapse, the primary bubble undergoes several decaying oscillations, producing a sequence of pressure pulses that continue to propagate into the surrounding medium. In the far field, the radiated pressure can be approximated using the secondary Bjerknes force. (*16*)

$$p(r,t)\approx p_\infty+\rho\left(\frac{R\ddot{R}}{r}+\frac{2\dot{R}^2}{r}\right), \tag{S9}$$

which serves as the effective driving pressure acting on nearby MOF particles.

For the MOF nanotransducers, the confined gas within the pore channels can be approximated as a driven local oscillator subjected to the external pressure field $p(r, t)$. Its response amplitude is governed by several key factors, including the initial gas content or gas volume fraction within the MOF pores; the pore size, geometry, and connectivity, which impose confinement; the stiffness of the MOF framework and impedance mismatch with the surrounding medium; and the amplitude, duration, and spectral content of the incident pressure pulses generated by the primary cavitation bubble. Variations in strain amplification observed across different MOF systems (e.g., ZIF-8, ZIF-67, NU-1003) can therefore be interpreted as the result of differences in pore architecture, gas storage capacity, and mechanical properties of the framework, which collectively govern the efficiency of this confined, driven oscillatory response.

When the radiated pressure field interacts with gas-loaded MOF particles, the confined or adsorbed gas undergoes cyclic compression and expansion. Due to geometric confinement and interfacial constraints, these oscillations manifest as restricted gas-bubble oscillations, which efficiently convert incident acoustic energy into highly localized solid–liquid interfacial mechanical perturbations. This mechanism leads to enhanced and spatially concentrated strain fields in the vicinity of the particles. In contrast, blank nanoparticles lacking gas loading do not possess a compressible phase capable of storing and releasing mechanical energy. As a result, they exhibit only weak, passive perturbations without significant strain amplification or temporal persistence. This coupled process explains the sustained oscillatory strain response observed over the 256-μs experimental window, as well as the pronounced mechanical amplification associated with gas-loaded MOF nanotransducers.

**Animal experiments.**

All animal procedures were conducted in accordance with the National Institutes of Health Guide for the Care and Use of Laboratory Animals and were approved by the Institutional Animal Care and Use Committee (IACUC) at the University of Texas at Austin (AUP-2021-00086, AUP-2021-00162, and AUP-2024-00150). The Animal Resources Center at the University of Texas at Austin supported all experiments. Surgical instruments were sterilized before all procedures. Mice and rats were housed under a 12 h light–dark cycle with ad libitum access to food and water. The animal housing facility was maintained at 21–24 °C with 40–60% relative humidity.

***In vivo* ultrasound imaging**.

The *in vivo* performance of liquid-phase gas nanotransducers as ultrasound imaging contrast agents was evaluated in C57BL/6 mice (20–26 g, 12–16 weeks old; Jackson Laboratory). For imaging of subcutaneously injected ZIF-8 liquid-phase nanotransducers, mice were anesthetized with 1.5% isoflurane delivered via an anesthesia system (Vaporizer Sales & Service Inc.) and placed on a temperature-controlled heating pad maintained at 37 °C. Ophthalmic ointment was applied to protect the eyes during anesthesia. The hair over the lower abdominal region was shaved prior to subcutaneous injection of 100 µL of ZIF-8 liquid-phase nanotransducers (10 mg/mL), saline, or commercial microbubbles (Lumason®, $1 \times 10^6$ microbubbles/mL; kindly provided by Dr. Pengfei Song's laboratory). Approximately 5 mL of ultrasound gel (Aquasonic® 100) was then applied to the injection site. Ultrasound imaging was performed using a linear array transducer (Vevo® F2, LAZR-X system, FUJIFILM VisualSonics) coupled to the gel, and B-mode images were acquired for contrast evaluation.

Intravenous administration of ZIF-8 liquid-phase nanotransducers for kidney ultrasound imaging was performed in C57BL/6 mice using the same ultrasound imaging system (Vevo® F2, LAZR-X, FUJIFILM VisualSonics) in B-mode. Briefly, mice were anesthetized with isoflurane and placed in a lateral position on a temperature-controlled heating pad. Hair over the abdominal region was shaved, and anesthesia was maintained throughout the procedure. Physiological parameters, including respiratory rate and heart rate, were continuously monitored using the built-in monitoring system. The ultrasound transducer was positioned over the abdomen and adjusted to obtain a clear longitudinal view of the kidney. Subsequently, 100 µL of ZIF-8 liquid-phase nanotransducers (15 mg/mL), saline, or commercial microbubbles ($1 \times 10^6$ microbubbles/mL) were intravenously injected, and real-time ultrasound images were acquired in B-mode.

Intracranial injection of ZIF-8 liquid-phase nanotransducers was performed for long-term brain ultrasound imaging and biostability evaluation. Mice were anesthetized with 2.5% isoflurane using an anesthesia system (Vaporizer Sales & Service Inc.), and the head was secured in a stereotaxic frame (Kopf Instruments). Each animal received subcutaneous injections of meloxicam (5 mg/kg) and Ethiqa (3.25 mg/kg) for analgesia, and ophthalmic ointment was applied to protect the eyes prior to surgery. A total volume of 1200 nL of ZIF-8 liquid-phase nanotransducers (200 µg/mL), saline, or blank ZIF-8 nanoparticles (gas-free, 200 µg/mL) was unilaterally injected into the ventral tegmental area (VTA) using the following stereotaxic coordinates relative to bregma:

anteroposterior (AP) −3.08 mm, mediolateral (ML) +0.40 mm, and dorsoventral (DV) −5.0 mm. After a recovery period of several days, mice were used for ultrasound imaging. Animals were anesthetized, placed on a heated pad, and maintained under isoflurane anesthesia. Hair over the cranial region was shaved, and ultrasound gel was applied to the scalp. The transducer was positioned above the injection site, and B-mode ultrasound images were acquired at different time intervals to assess signal persistence and biostability.

***In vitro neural modulation mediated by liquid-phase nanotransducers and quantified via calcium imaging.***

Primary mouse cortical neurons were prepared as previously described.(*21*) Following isolation, neurons were plated in 24-well plates and cultured for 3 days. For calcium indicator expression, 1 μL of pAAV-hSyn-GCaMP6s-WPRE-SV40 (titer ≥ 1 × $10^{13}$ vg/mL) was added to each well. After 4–6 days of viral transduction, liquid-phase ZIF-8 nanotransducers were introduced into the culture medium at a final concentration of 50 μg/mL. Neurons were then imaged on a fluorescence microscope, with an ultrasound transducer immersed in the culture medium. Ultrasound stimulation (primary peak pressure: 0.19 MPa, frequency: 1.5 MHz, pulse scheme: 300 μs on and 700 μs off, total duration 1 s) was applied to activate the nanotransducers and induce neuronal activation. Time-lapse fluorescence videos were acquired using a Leica DMi8 fluorescence microscope equipped with a 20× air objective, with a 5 ms exposure time in the green channel (excitation: 450–490 nm). Calcium responses were quantified by calculating the normalized fluorescence change (ΔF/F). Fluorescence time-series data were extracted from 100 neurons per condition by manual segmentation, followed by grayscale conversion using ImageJ. The raw fluorescence traces were further processed using a custom MATLAB algorithm, which detrended and normalized the data via second-order polynomial fitting and baseline maximum fluorescence extraction to compensate for photobleaching. Neurons treated with blank ZIF-8 nanoparticles (without gas loading) under identical ultrasound parameters served as a negative control.

***Fiber photometry measurements in the VTA during liquid-phase gas nanotransducer -mediated ultrasound stimulation.***

To enable real-time evaluation of neural activation, GCaMP6s-transduced mice were generated. C57BL/6 mice (20–26 g; 8–12 weeks old; Jackson Laboratory) were used in all experiments. Mice were anesthetized with 2.5% isoflurane using an anesthesia machine (Vaporizer Sales & Service Inc.), and the head was secured in a stereotaxic frame (Kopf Stereotaxic Instruments). Each animal received a subcutaneous injection of meloxicam (5 mg/kg) and Ethiqa (3.25 mg/kg), and ophthalmic ointment was applied to prevent corneal drying. For fiber photometry experiments, 1200 nL of pAAV-hSyn-GCaMP6s-WPRE-SV40 (titer $\geq 1 \times 10^{13}$ vg/mL) was unilaterally injected into the ventral tegmental area (VTA) using the following stereotaxic coordinates relative to bregma: anteroposterior (AP) −3.08 mm, mediolateral (ML) +0.40 mm, and dorsoventral (DV) −5.0 mm. Mice were allowed to recover for four weeks to ensure stable GCaMP6s expression before subsequent experiments.

For sonogenetic stimulation, 2 µL of liquid-phase gas ZIF-8 nanotransducers (200 µg/mL) were unilaterally injected into the VTA using the same stereotaxic coordinates as the viral injection. An optical fiber was then implanted above the virus-transduced VTA region (AP −3.08 mm, ML +0.50 mm, DV −5.0 mm). After recovery, mice were head-fixed, and focused ultrasound ( primary peak pressure 0.50 MPa, frequency 1.5 MHz, pulse scheme 300 µs on and 700 µs off, total duration 3 s) was applied to the VTA region. The FUS focal length was set to 5 mm by adjusting the water coupling distance to cover the VTA region. Calcium signals were recorded using an R820 Dual-Color Multi-Channel Fiber Photometry System (RWD Life Science) throughout the sonogenetic stimulation protocol at different time intervals.

***c-Fos immunofluorescence staining and histological assessment of mouse brain sections.***

Mice that had received unilateral injections of liquid-phase gas ZIF-8 nanotransducers into the VTA and were maintained for 60 days were first subjected to ultrasound stimulation. 60 minutes after ultrasound treatment, mice were deeply anesthetized by intraperitoneal injection of ketamine (16 mg/kg). Transcardial perfusion was then performed with phosphate-buffered saline (PBS), followed by 4% paraformaldehyde (PFA). Brains were extracted, post-fixed in 4% PFA at 4 °C overnight, and sectioned into 50 µm slices using a vibrating blade microtome (Leica VT1200). Brain sections were washed with PBS containing 0.3% Triton X-100 and subsequently blocked with 5% bovine serum albumin (BSA) in PBS for 30 min at room temperature. After blocking, sections were incubated with primary antibodies diluted in PBS containing 0.3% Triton X-100: rabbit anti-c-Fos (ab222699, Abcam, 1:500) and mouse anti-tyrosine hydroxylase (MA1-24654,

Fisher Scientific, 1:1000). Samples were incubated overnight at 4 °C and washed three times with PBS. Sections were then incubated with secondary antibodies diluted in PBS, including goat anti-rabbit Alexa Fluor 405 (ab175652, Abcam, 1:500), goat anti-mouse Alexa Fluor 488 (ab150113, Abcam, 1:1000), and Hoechst 33342 (17535, ATT Bioquest, 1:5000), for 2 h at room temperature in the dark. After three washes with PBS, slices were mounted using mounting medium (9990402, Fisher Scientific) and covered with a coverslip. Confocal images were acquired using a Zeiss 710 laser scanning confocal microscope. Detailed antibody information is provided in **Supplementary Table 1**.

***Biosafety evaluation through Iba1, GFAP, and cleaved caspase-3 immunostaining.***

For histological evaluation of neuroinflammation and cytotoxicity, mice were sacrificed 14 days after in vivo sonogenetic procedures. Animals were anesthetized via intraperitoneal injection of ketamine (16 mg/kg), followed by transcardial perfusion with PBS and 4% PFA. Brains were extracted, post-fixed in 4% PFA at 4 °C overnight, and sectioned into 60 μm slices using a vibrating blade microtome. Brain sections were washed with PBS and blocked with 5% BSA in PBS for 30 min at room temperature. The blocking buffer was then replaced with one of the following primary antibody solutions diluted in PBS: rabbit anti-Iba1 (013-27691, Wako Chemicals, 1:500), rabbit anti-cleaved caspase-3 (9661, Cell Signaling Technology, 1:500), or rabbit anti-GFAP (13-0300, Invitrogen, 1:500). Sections were incubated overnight at 4 °C and washed three times with PBS. Subsequently, sections were incubated with donkey anti-rabbit Alexa Fluor 594 (A32754, Invitrogen, 1:500) and Hoechst 33342 (17535, ATT Bioquest, 1:5000) for 2 h at room temperature in the dark. After washing three times with PBS, slices were mounted using mounting medium and sealed with coverslips. Confocal images were obtained using a Zeiss 710 laser scanning confocal microscope. Antibody details are listed in **Supplementary Table 1**.

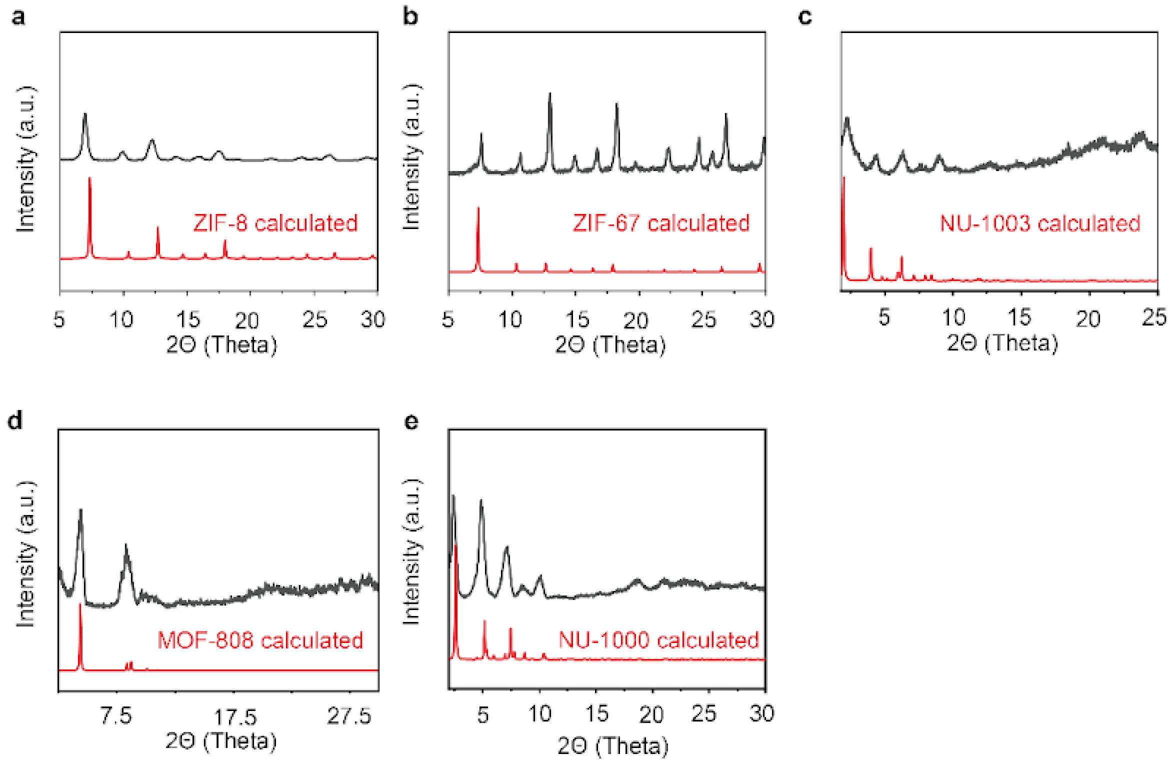


**Supplementary Fig.1**. The X-ray diffraction (XRD) tests of MOF nanocrystals, (a) ZIF-8, (b) ZIF-67, (c) NU-1003, (d) MOF-808, and (e) NU-1000.

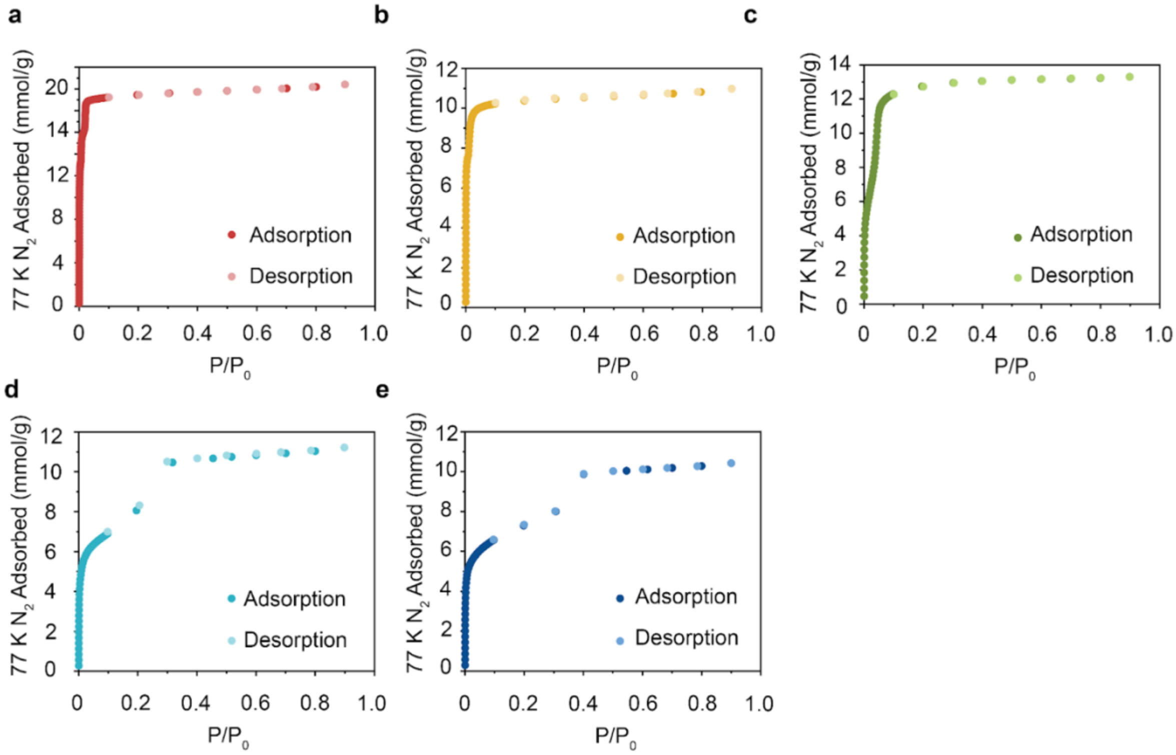


**Supplementary Fig.2**. The porosity characterization of different MOF nanocrystals through single-component sorption isotherms of nitrogen at 77 K (a) ZIF-8, (b) ZIF-67, (c) MOF-808, (d) NU-1000, and (e) NU-1003.

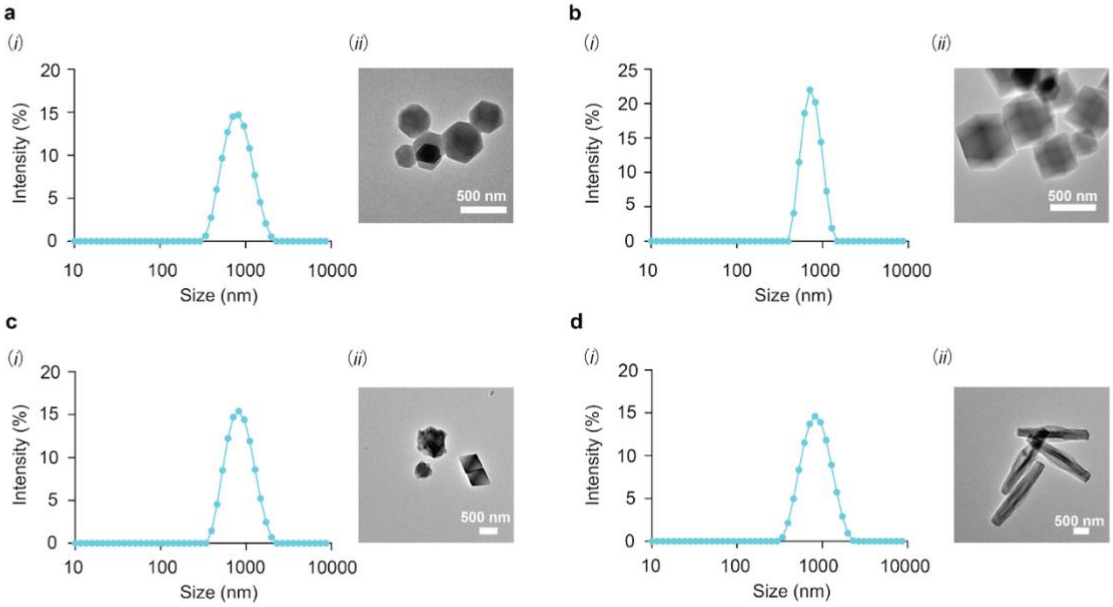


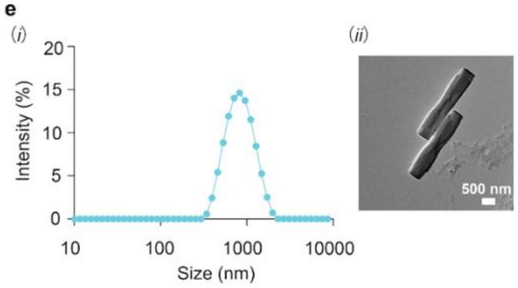


f

| Dynamic light scattering tests of MOFs | | |
|---|---|---|
| MOFs | size (nm) | PDI |
| ZIF-8 | 762.0 ± 22.9 | 0.345 |
| ZIF-67 | 737.7 ± 24.4 | 0.383 |
| MOF-808 | 795.1 ± 4.9 | 0.329 |
| NU-1000 | 828.3 ± 14.5 | 0.393 |
| NU-1003 | 774.9 ± 36.5 | 0.387 |

**Supplementary Fig.3**. Transmission Electron Microscope (TEM) images and hydrodynamic size distribution measured by DLS of the different MOF nanocrystals. (a) ZIF-8, (b) ZIF-67, (c) MOF-808, (d) NU-1000, (e) NU-1003, and (f) the details of the size distribution of all MOF nanocrystals.

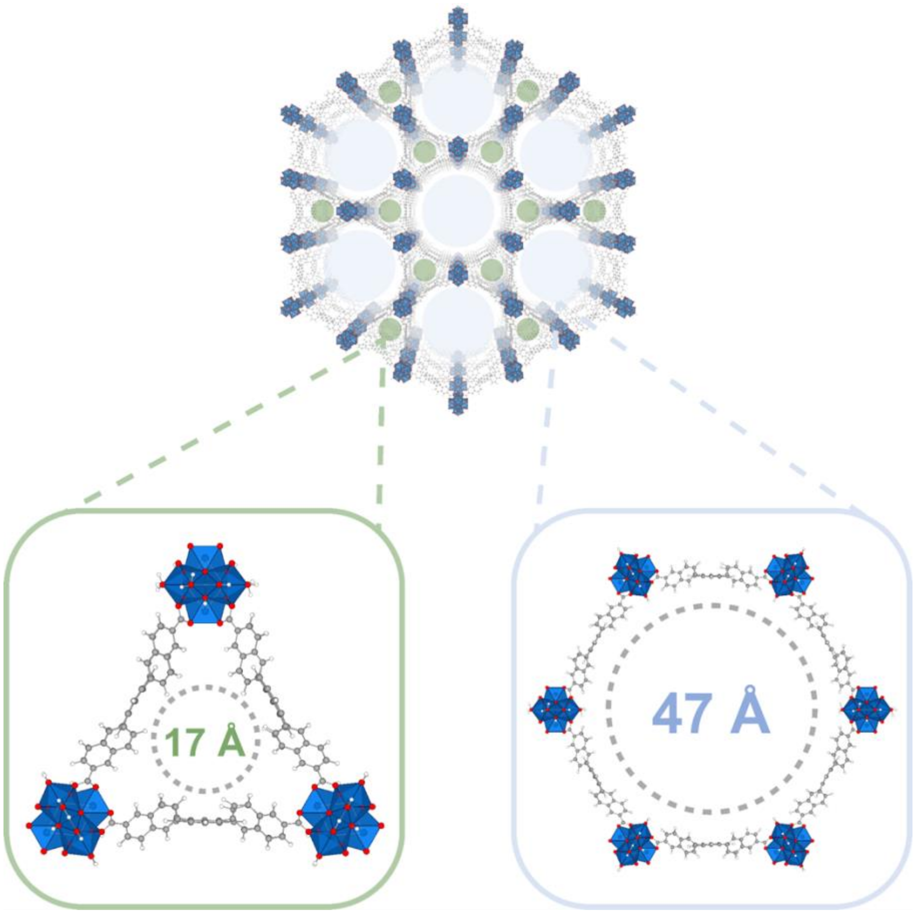


**Supplementary Fig.4**. Crystal structure of NU-1003 showing smaller triangular channels (green) and larger hexagonal channels (blue) with reported pore apertures.(*22*)

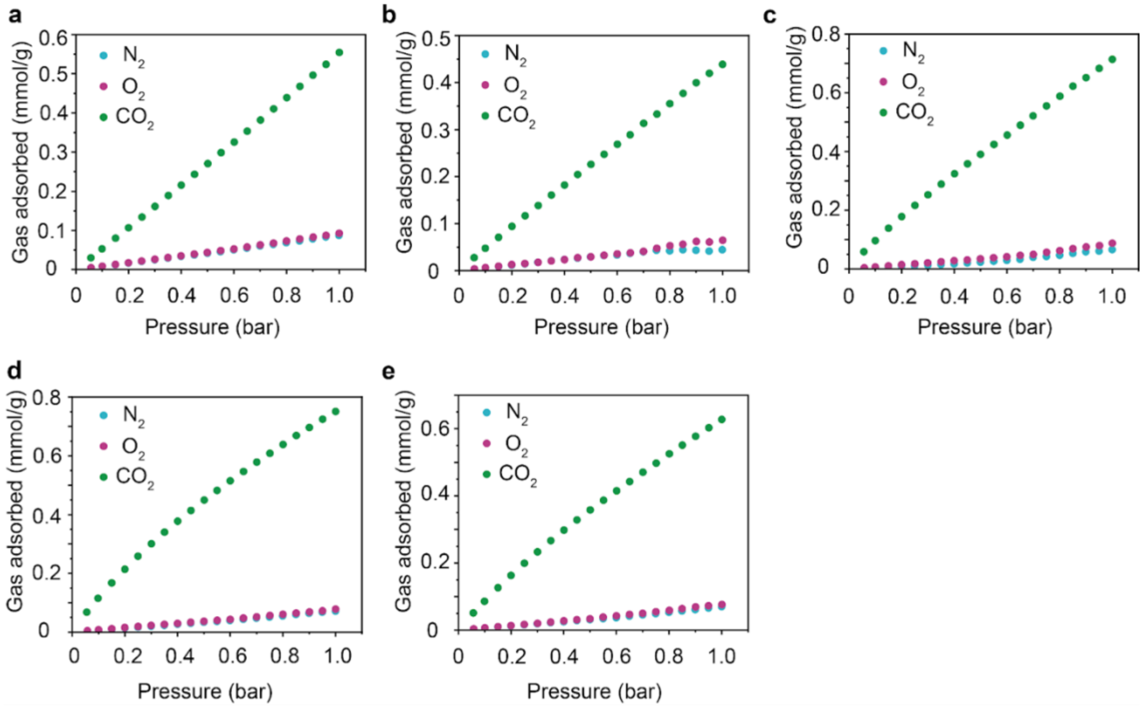


**Supplementary Fig.5**. Quantitative comparison of the loading capacities of different gases, including $CO_2$, $N_2$, $O_2$, and air in (a) ZIF-8, (b) ZIF-67, (c) MOF-808, (d) NU-1000 and (e) NU-1003 at 37 °C.

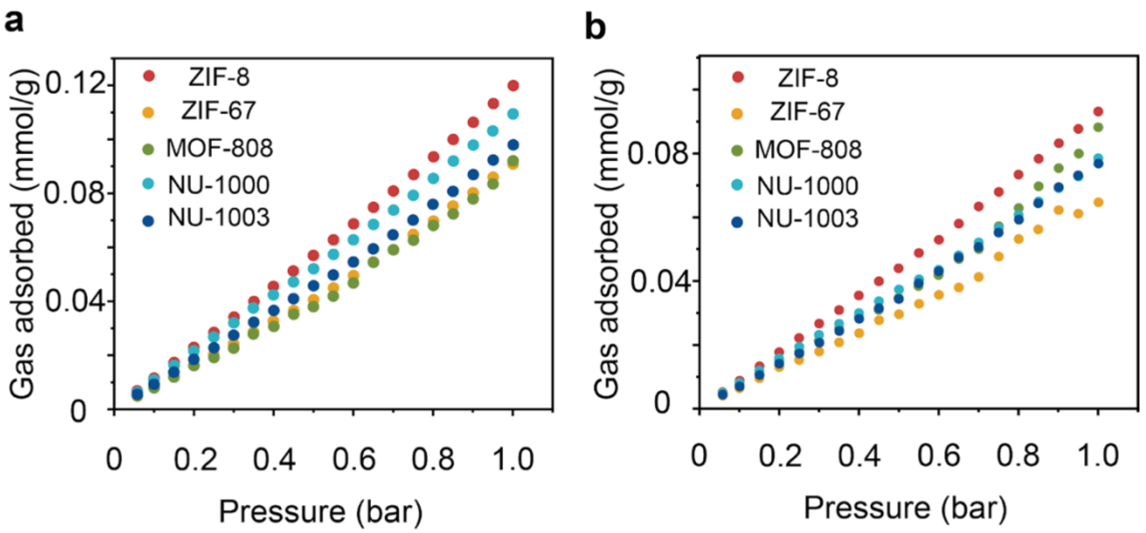


**Supplementary Fig.6**. Quantitative comparison of the loading capacities of $O_2$, and air in ZIF-8 at (a) 20 °C and (b) 37 °C.

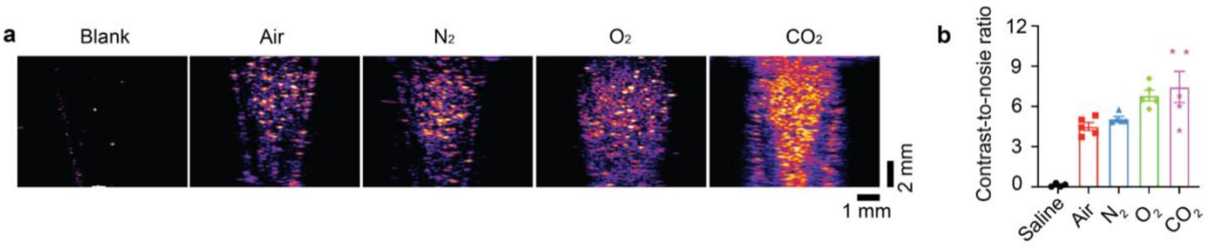


**Supplementary Fig.7**. (a) Ultrasound imaging (27 MHz) of ZIF-8 loaded with different designer gases, including air, $N_2$, $O_2$ and $CO_2$. (b) CNR quantification for ZIF-8 with different gas loadings.

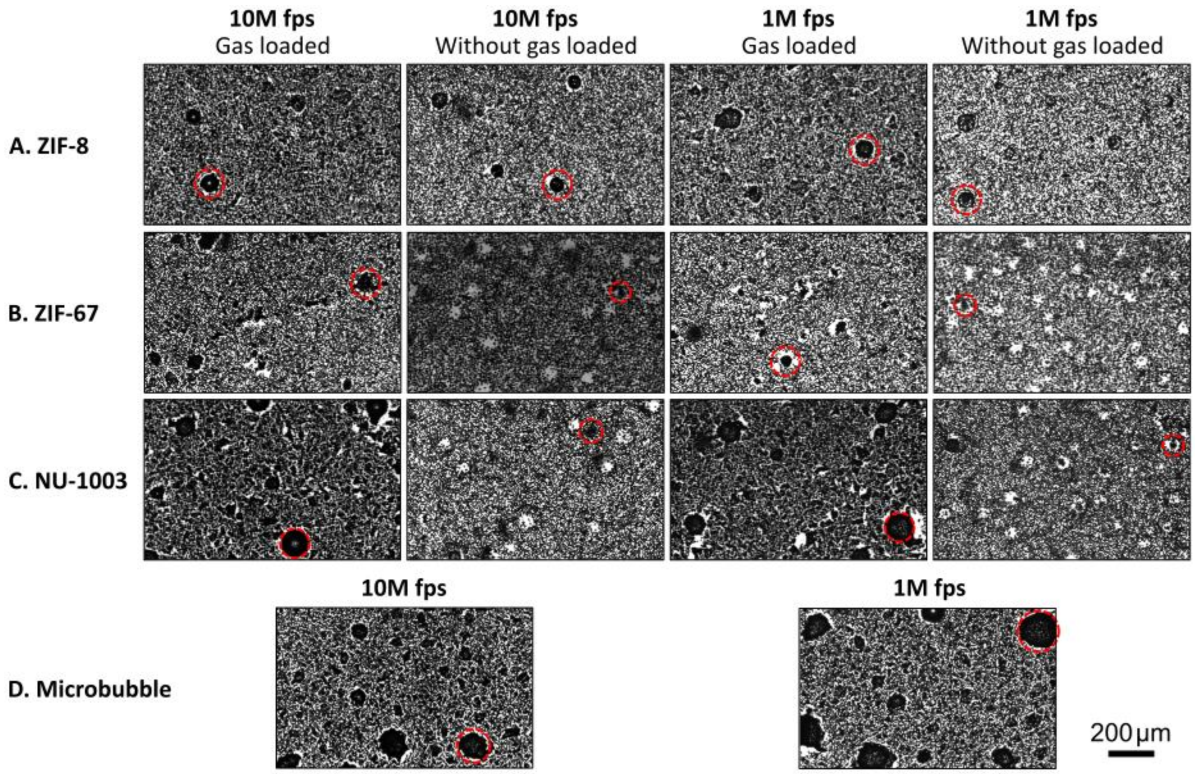


**Supplementary Fig.8**. Representative microscopy imaging of metal organic framework (MOF) particles and microbubbles embedded in gelation with DIC speckle patterns. Rows A–C show samples containing different MOF particles: (A) ZIF-8, (B) ZIF-67, and (C) NU-1003. For each MOF type, images are shown under both gas-loaded and unloaded conditions, captured at two different frame rates: 10 million fps and 1 million fps. Row D shows embedded microbubbles as a comparison case, also imaged at 10 million fps and 1 million fps. The dense background texture corresponds to the embedded toner speckle pattern used for DIC. Red dashed circles highlight representative MOF particles or microbubbles selected for laser-induced cavitation experiments. Gas-loaded MOF particles exhibit distinct optical contrast and bubble nucleation features compared to unloaded particles. Scale bar: 200 μm.

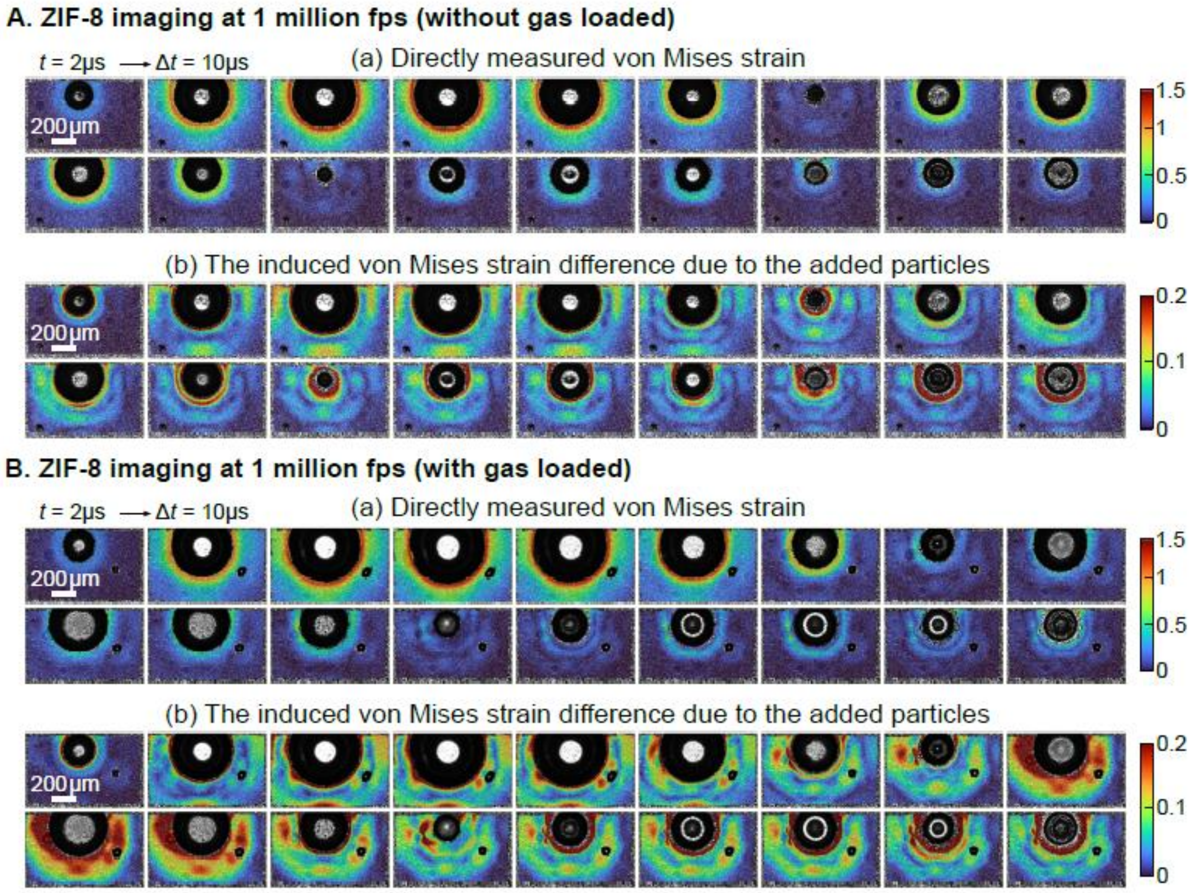


**Supplementary Fig.9**. Time-resolved full-field von Mises strain maps of ZIF-8 particles under laser-induced cavitation at 1 million fps. (A) ZIF-8 without gas loading. (a) Directly measured von Mises strain fields showing the evolution of the primary cavitation bubble and surrounding deformation at selected time points. (b) Induced von Mises strain difference highlighting the particle-induced strain contribution after subtracting the background bubble field. (B) ZIF-8 with gas loading. (a) Directly measured von Mises strain fields under identical loading conditions at selected time points. (b) Induced von Mises strain difference maps, revealing significantly enhanced strain localization and amplification around gas-loaded particles compared to the unloaded case.

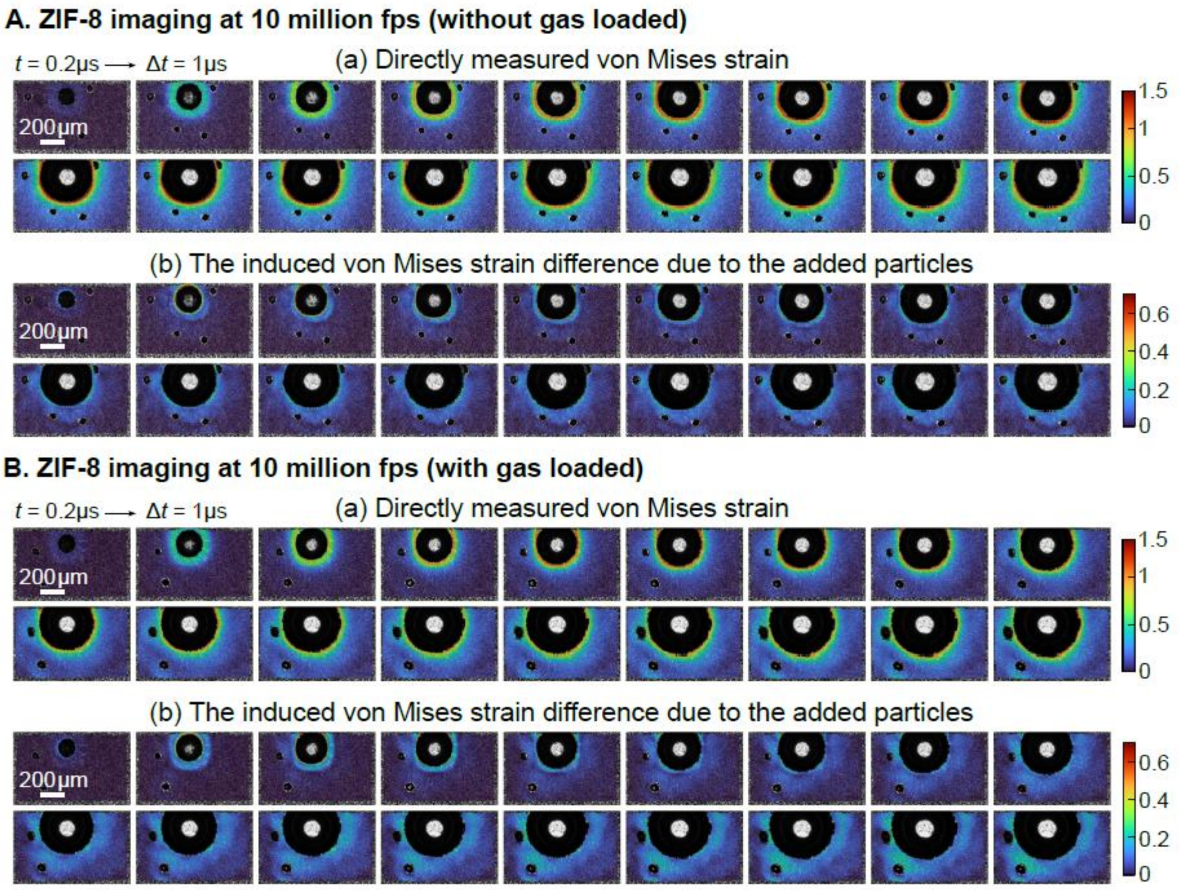


**Supplementary Fig.10**. Time-resolved full-field von Mises strain maps of ZIF-8 particles under laser-induced cavitation at 10 million fps. (A) ZIF-8 without gas loading. (a) Directly measured von Mises strain fields showing the evolution of the primary cavitation bubble and surrounding deformation at selected time points. (b) Induced von Mises strain difference highlighting the particle-induced strain contribution after subtracting the background bubble field. (B) ZIF-8 with gas loading. (a) Directly measured von Mises strain fields under identical loading conditions at selected time points. (b) Induced von Mises strain difference maps, revealing significantly enhanced strain localization and amplification around gas-loaded particles compared to the unloaded case.

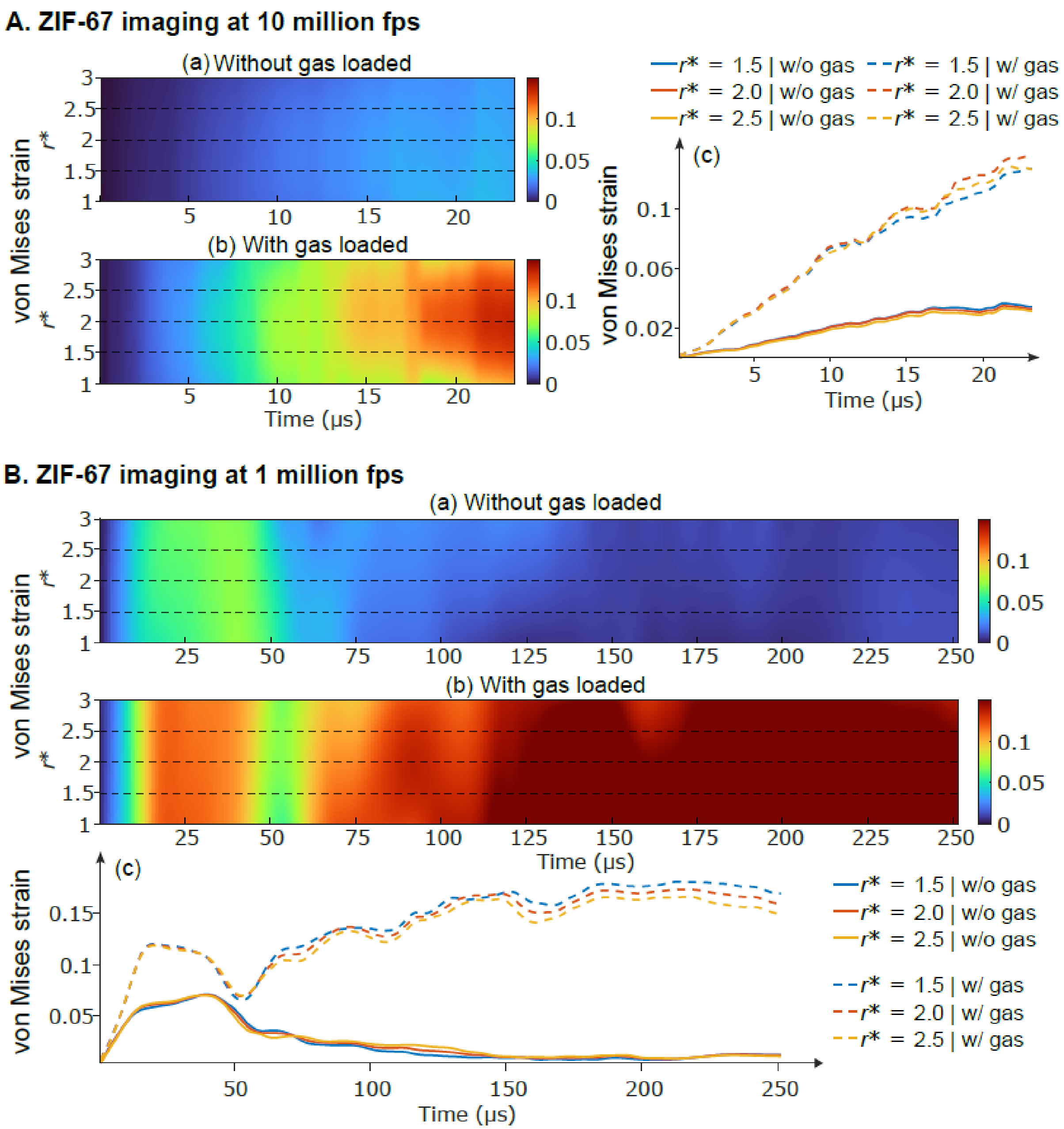


**Supplementary Fig.11**. Spatiotemporal evolution of particle-induced von Mises strain field around ZIF-67 particles under laser-induced cavitation at different imaging rates. (A) Results acquired at 10 million fps. Kymograph diagrams show the radial vs. temporal distribution of particle-induced von Mises strain as a function of normalized radial position $r^*$ for (a) without gas loading and (b) with gas loading. (c) Corresponding radial strain histories extracted at $r^* = 1.5$, 2.0, and 2.5, comparing gas-loaded and unloaded conditions. Gas-loaded ZIF-67 particles exhibit significantly amplified and sustained strain fields, particularly at larger radial positions. (B) Results acquired at 1 million fps: (a–b) Kymograph diagrams of particle-induced von Mises strain field around unloaded and gas-loaded particles, respectively. (c) Extracted strain histories at selected $r^*$ values, showing consistent strain enhancement under gas-loaded conditions, though with reduced temporal resolution compared to 10 million fps imaging.

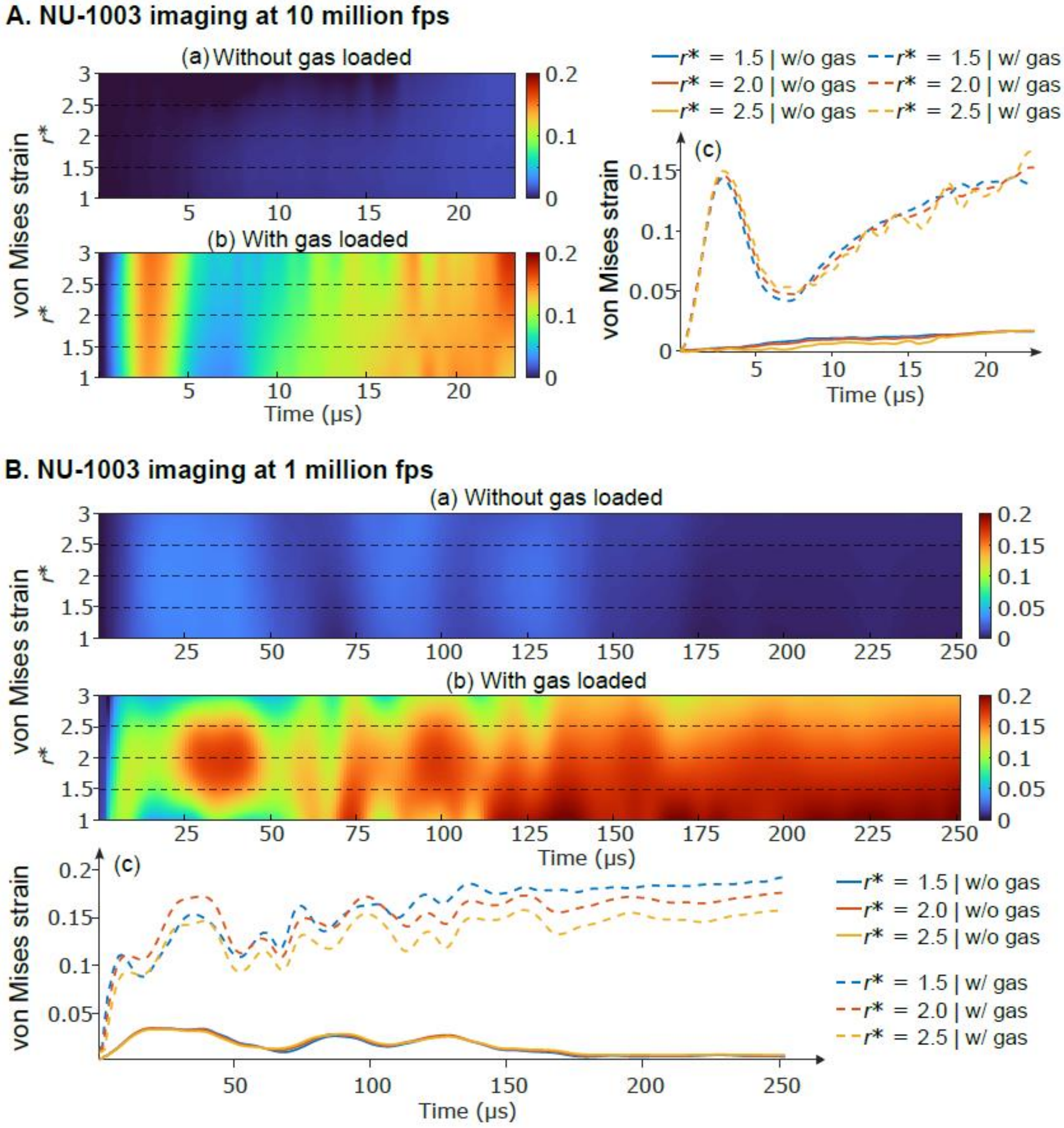


**Supplementary Fig.12**. Spatiotemporal evolution of particle-induced von Mises strain around NU-1003 particles under laser-induced cavitation at different imaging rates. (A) Results acquired at 10 million fps. Kymograph diagrams show the radial vs. temporal distribution of particle-induced von Mises strain as a function of normalized radial position $r^*$ for (a) without gas loading and (b) with gas loading. (c) Corresponding radial strain histories extracted at $r^*$ = 1.5, 2.0, and 2.5, comparing gas-loaded and unloaded conditions. Gas-loaded NU-1003 particles exhibit significantly amplified and sustained strain fields, particularly at larger radial positions. (B) Results acquired at 1 million fps. (a–b) Kymograph diagrams of particle-induced von Mises strain field aroundunloaded and gas-loaded particles, respectively. (c) Extracted strain histories at selected $r^*$ values, showing consistent strain enhancement under gas-loaded conditions, though with reduced temporal resolution compared to 10 million fps imaging.

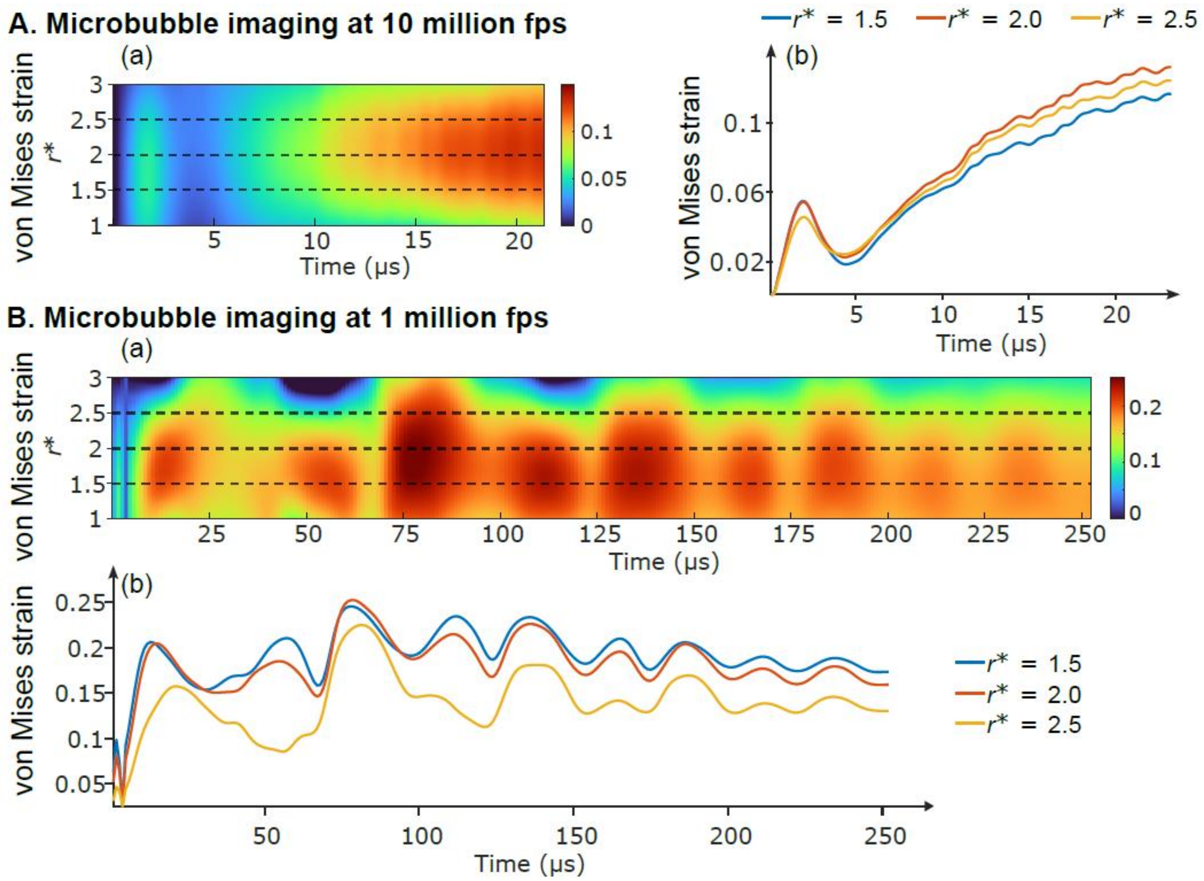


**Supplementary Fig.13**. Spatiotemporal evolution of particle-induced von Mises strain field around microbubbles under laser-induced cavitation at different imaging rates. (A) Results acquired at 10 million fps. (a) Kymograph diagrams show the radial vs. temporal distribution of particle-induced von Mises strain as a function of normalized radial position $r*$. (b) Corresponding radial strain histories extracted at $r^*$ = 1.5, 2.0, and 2.5. (B) Results acquired at 1 million fps. (a)

Kymograph diagrams of particle-induced von Mises strain field around microbubbles. (b) Extracted strain histories at selected $r^*$ values, showing consistent strain enhancement under gas-loaded conditions, though with reduced temporal resolution compared to 10 Mfps imaging.

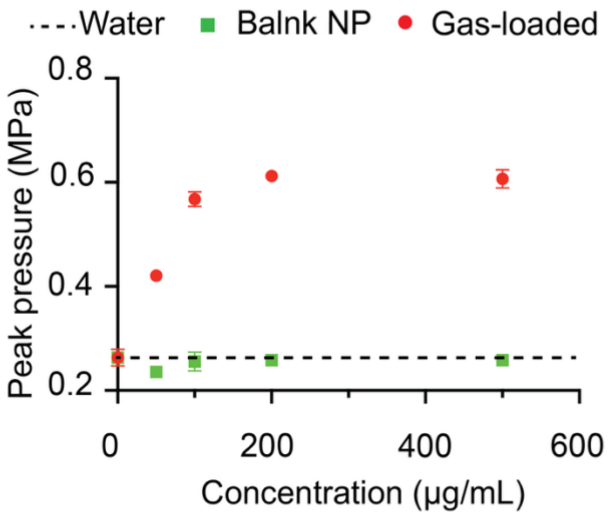


**Supplementary Fig.14**. Acoustic peak pressure of ZIF-8 nanotransducers with or without gas loading measured across concentrations under 1.5 MHz focused ultrasound at identical output power.

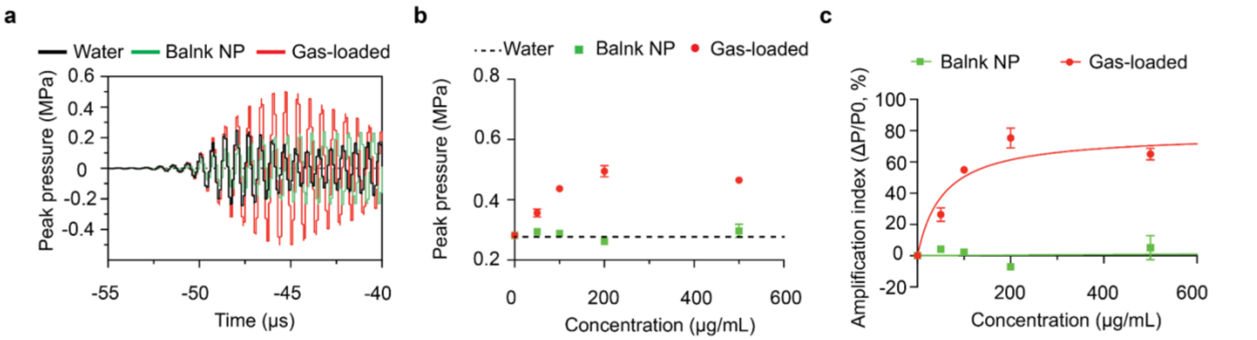


**Supplementary Fig.15**. Gas-bubble oscillations in ZIF-67 liquid-phase nanotransducers enable acoustic actuation. (a) Acoustic waveforms recorded in aqueous solutions containing ZIF-67 nanotransducers (200 μg mL$^{-1}$) with or without gas loading, and in bulk water, under 1.5 MHz focused ultrasound (FUS) at identical output power. (b) Peak acoustic pressure of ZIF-67 nanotransducers, with or without gas loading, as a function of concentration under 1.5 MHz FUS at identical output power. (c) Acoustic actuation of ZIF-67 nanotransducers, with or without gas loading, across concentrations under 1.5 MHz FUS at identical output power.

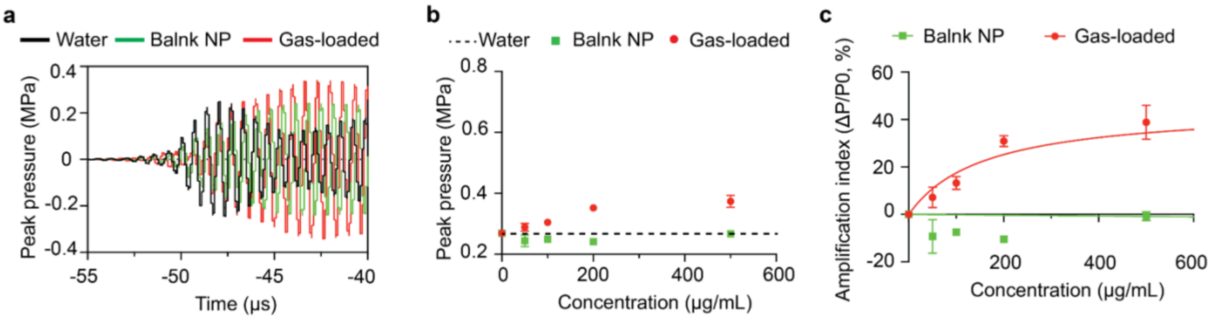


**Supplementary Fig.16**. Gas-bubble oscillations in NU-1003 liquid-phase nanotransducers enable acoustic actuation. (a) Acoustic waveforms recorded in aqueous solutions containing NU-1003 nanotransducers (200 μg mL$^{-1}$) with or without gas loading, and in bulk water, under 1.5 MHz focused ultrasound (FUS) at identical output power. (b) Peak acoustic pressure of NU-1003 nanotransducers, with or without gas loading, as a function of concentration under 1.5 MHz FUS at identical output power. (c) Acoustic actuation of NU-1003 nanotransducers, with or without gas loading, across concentrations under 1.5 MHz FUS at identical output power.

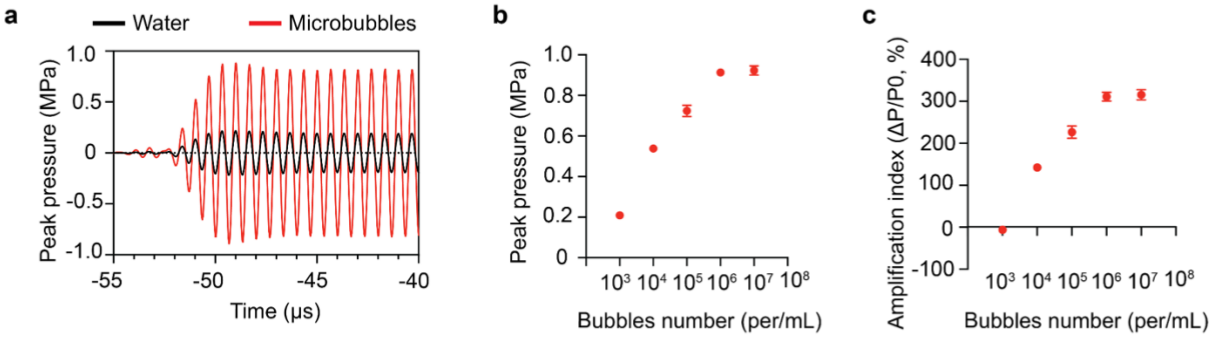


**Supplementary Fig.17**. Microbubble oscillations in aqueous solution enable acoustic actuation. (a) Acoustic waveforms recorded in aqueous solutions containing microbubbles ($10^7$ microbubbles mL$^{-1}$) in bulk water, under 1.5 MHz focused ultrasound (FUS) at identical output power. (b) Peak acoustic pressure of microbubbles, as a function of concentration under 1.5 MHz FUS at identical output power. (c) Acoustic actuation of microbubbles, across concentrations under 1.5 MHz FUS at identical output power.

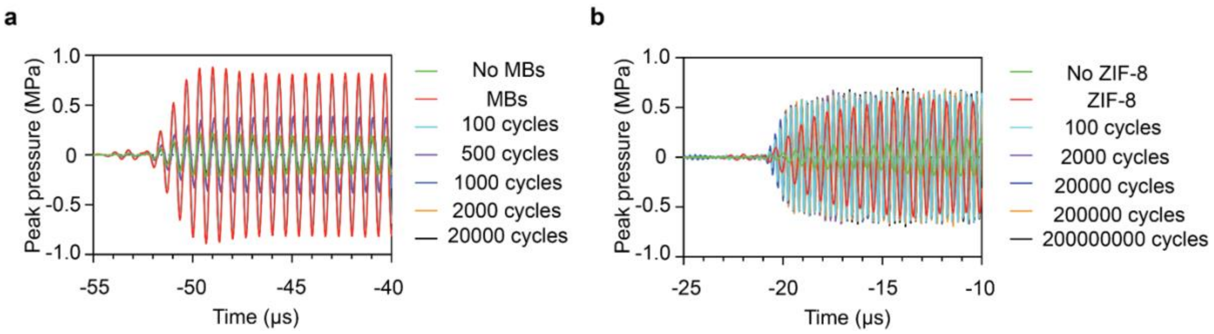


**Supplementary Fig.18**. **Acoustic stability under repeated ultrasound stimulation.** (a) Acoustic waveforms recorded in aqueous solutions containing microbubbles ($10^7$ microbubbles $mL^{-1}$) after different numbers of ultrasound cycles (300 µs on, 700 µs off; PRF 1,000 Hz; peak pressure 0.3 MPa) under 1.5 MHz focused ultrasound (FUS) at identical output power. (b) Acoustic waveforms recorded in aqueous solutions containing ZIF-8 liquid-phase gas nanotransducers (200 µg $mL^{-1}$) after different numbers of ultrasound cycles under the same conditions.

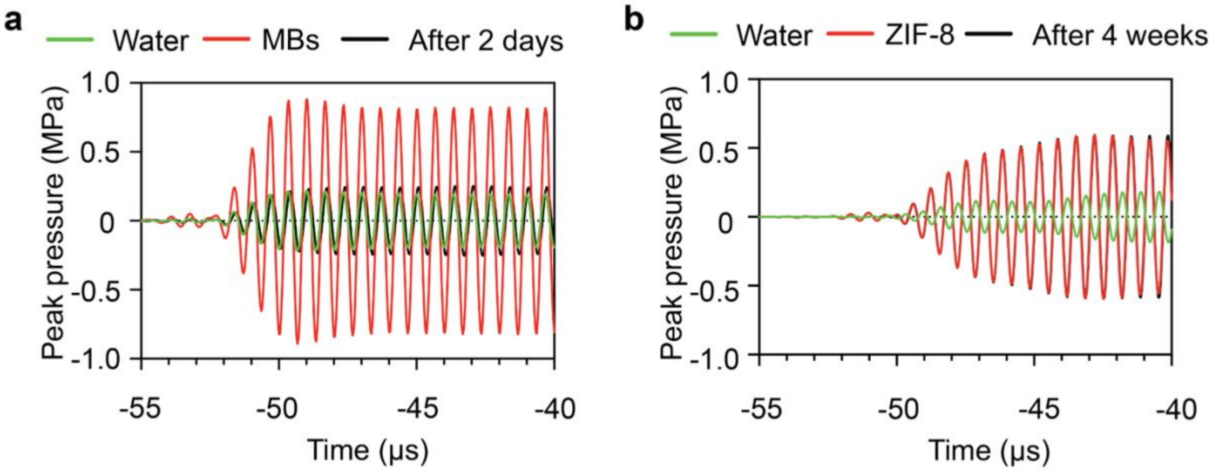


**Supplementary Fig. 19 Acoustic stability during storage at 37 °C.** (a) Acoustic waveforms recorded from aqueous solutions containing microbubbles ($10^7$ $mL^{-1}$) after incubation at 37 °C for 2 days. (b) Acoustic waveforms recorded from aqueous solutions containing ZIF-8 liquid-phase gas nanotransducers (200 µg $mL^{-1}$) after incubation at 37 °C for 4 weeks.

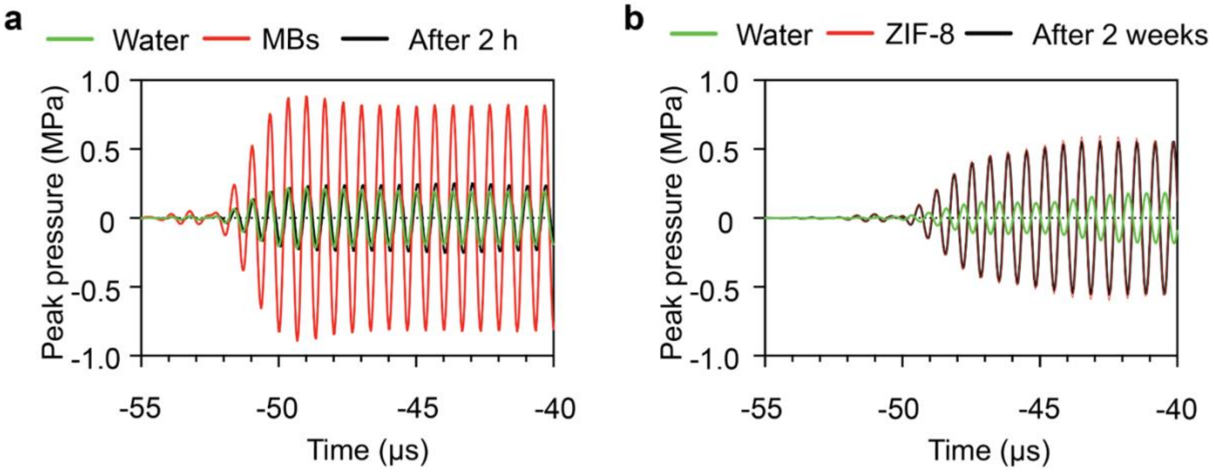


**Supplementary Fig. 20 Acoustic stability during storage at 60 °C.** (a) Acoustic waveforms recorded from aqueous solutions containing microbubbles ($10^7$ $mL^{-1}$) after incubation at 60 °C for 2 h. (b) Acoustic waveforms recorded from aqueous solutions containing ZIF-8 liquid-phase gas nanotransducers (200 μg $mL^{-1}$) after incubation at 60 °C for 2 weeks.

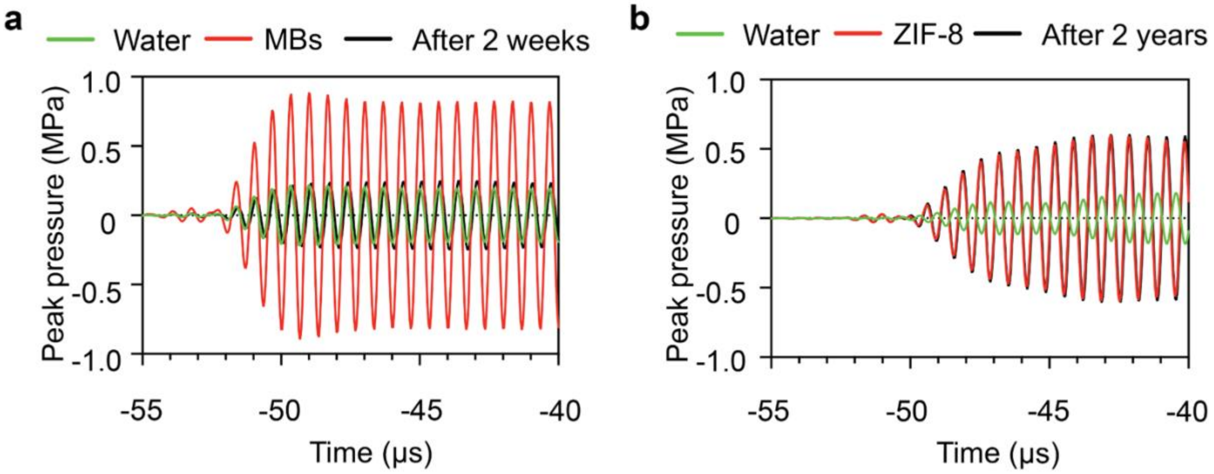


**Supplementary Fig. 21 Acoustic stability during storage at room temperature.** (a) Acoustic waveforms recorded from aqueous solutions containing microbubbles ($10^7$ $mL^{-1}$) after incubation at room temperature for 2 weeks. (b) Acoustic waveforms recorded from aqueous solutions containing ZIF-8 liquid-phase gas nanotransducers (200 μg $mL^{-1}$) following storage at room temperature for 2 years.

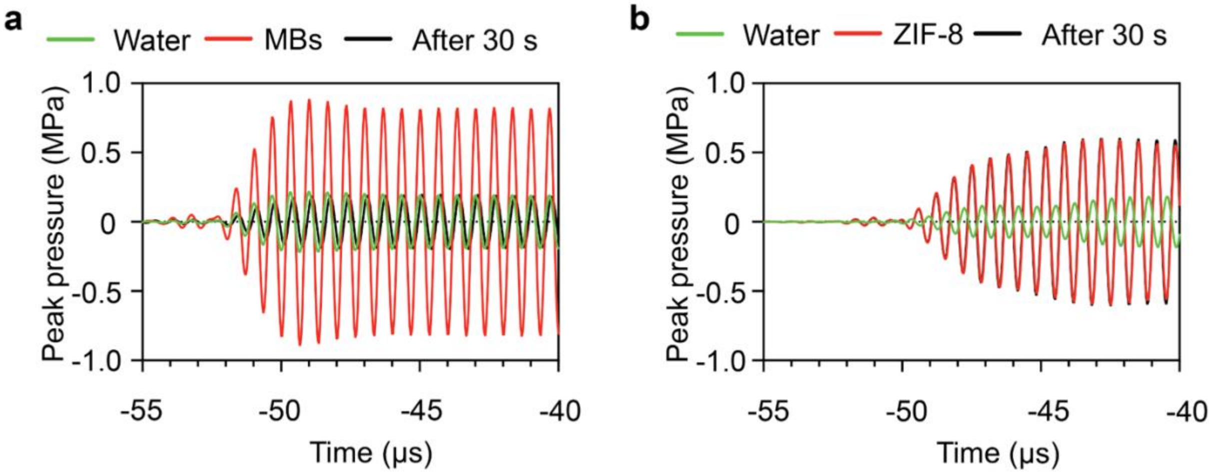


**Supplementary Fig. 22. Acoustic stability during storage at 10 MPa peak pressure.** (a) Acoustic waveforms recorded from aqueous solutions containing microbubbles ($10^7$ $mL^{-1}$) after incubation at 10 MPa peak pressure for 30 s. (b) Acoustic waveforms recorded from aqueous solutions containing ZIF-8 liquid-phase gas nanotransducers (200 μg $mL^{-1}$) after incubation at 10 MPa peak pressure for 30 s.

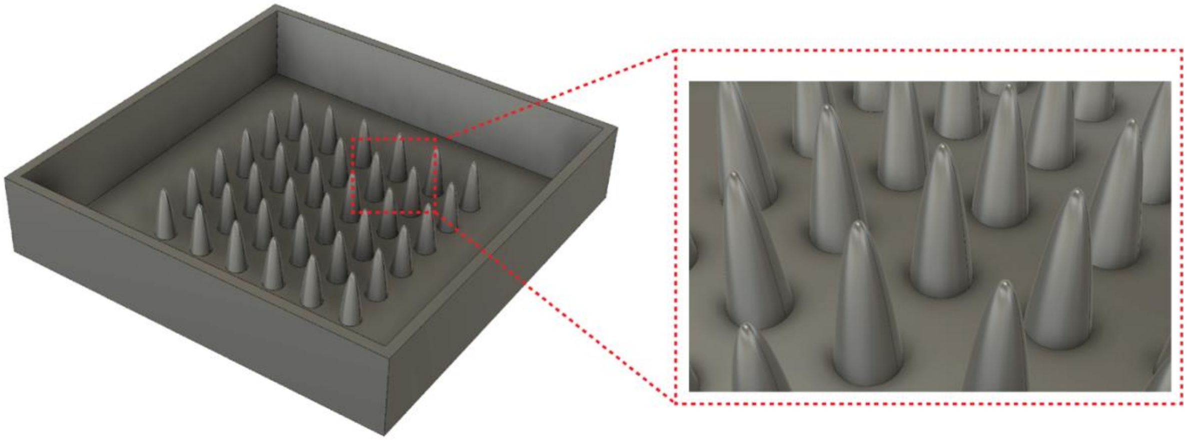

**Supplementary Fig. 23.** The mold was first designed to create the silicone phantom structure.

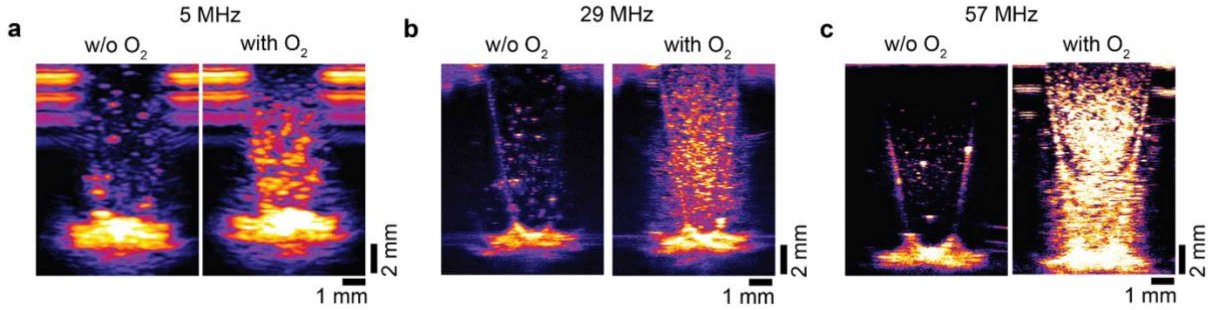


**Supplementary Fig. 24. Frequency-dependent ultrasound imaging performance.** Ultrasound images of a silicone phantom containing non-gas loaded ZIF-8 nanocrystals (Blank NP) or ZIF-8 liquid-phase gas nanotransducers (2 mg $mL^{-1}$, ~800 nm) acquired at different frequencies: (a) 5 MHz, (b) 29 MHz and (c) 57 MHz.

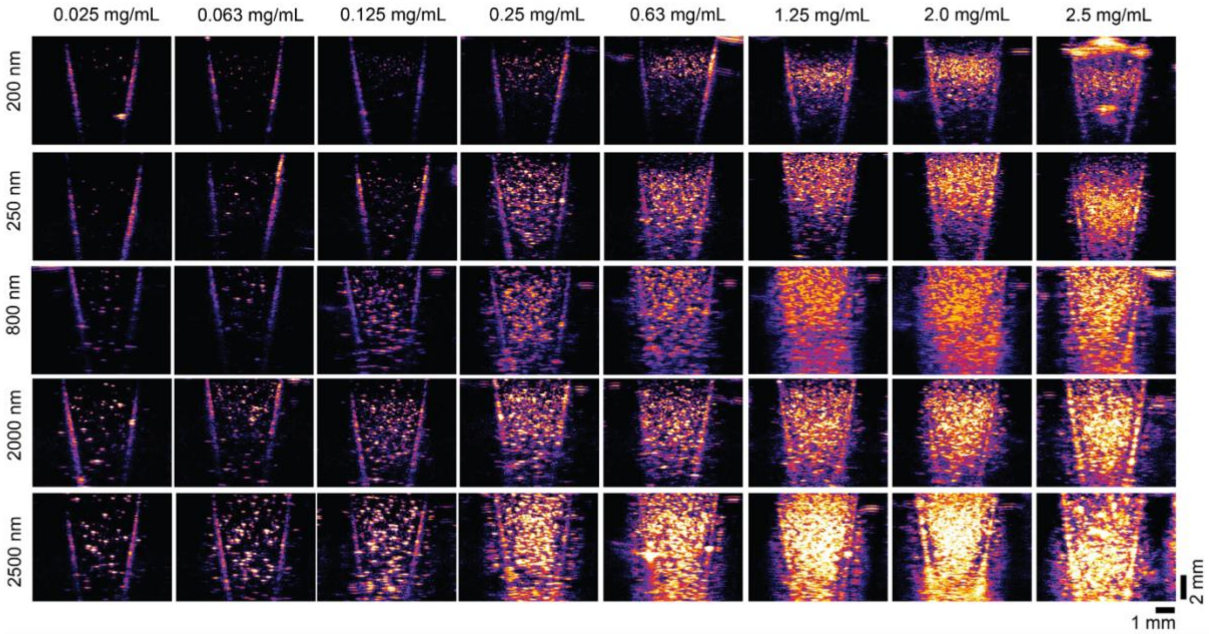


**Supplementary Fig. 25. Ultrasound imaging performance at 57 MHz.** Ultrasound images of a silicone phantom containing ZIF-8 liquid-phase gas nanotransducers acquired at varying concentrations and hydrodynamic sizes.

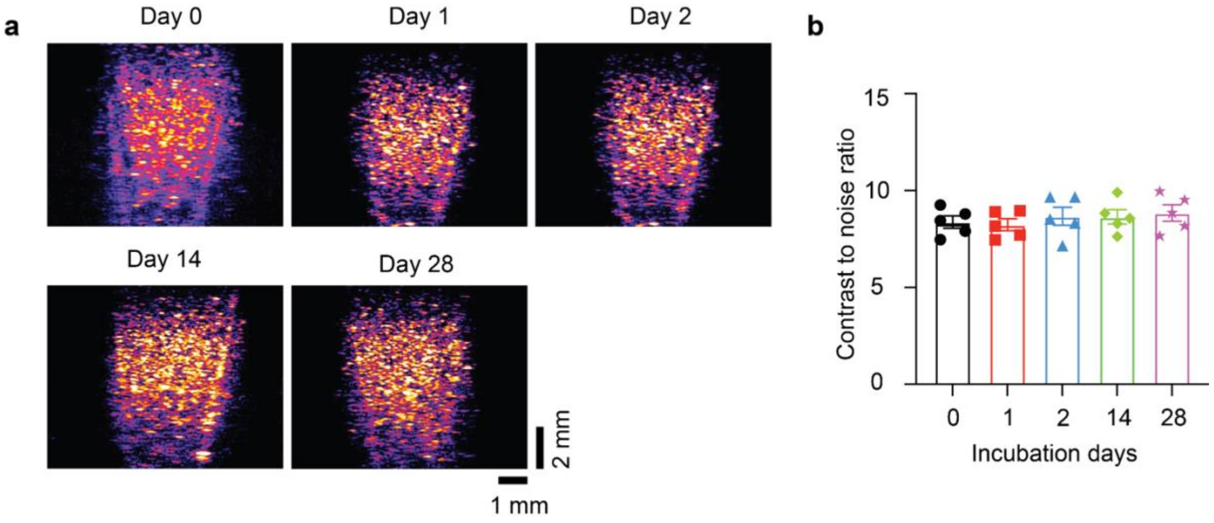


**Supplementary Fig. 26 Ultrasound imaging stability of ZIF-8 liquid-phase gas nanotransducers under physiological incubation conditions.** (a) Ultrasound images of silicone phantoms containing ZIF-8 liquid-phase gas nanotransducers acquired at 57 MHz after incubation at 37 °C for the indicated durations.(b) Quantification of the CNR of ZIF-8 liquid-phase gas nanotransducers following incubation at 37 °C.

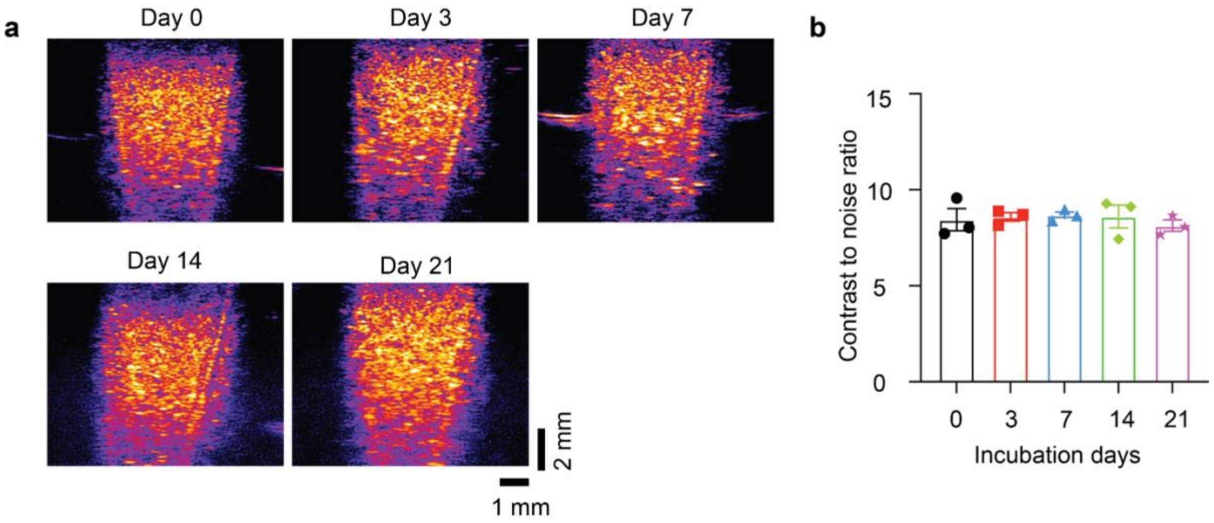


**Supplementary Fig. 27. Ultrasound imaging stability of ZIF-8 liquid-phase gas nanotransducers under elevated-temperature conditions.** (a) Ultrasound images of silicone phantoms containing ZIF-8 liquid-phase gas nanotransducers acquired at 57 MHz after incubation at 60 °C for the indicated durations. (b) Quantification of the CNR of ZIF-8 liquid-phase gas nanotransducers following incubation at 60 °C.

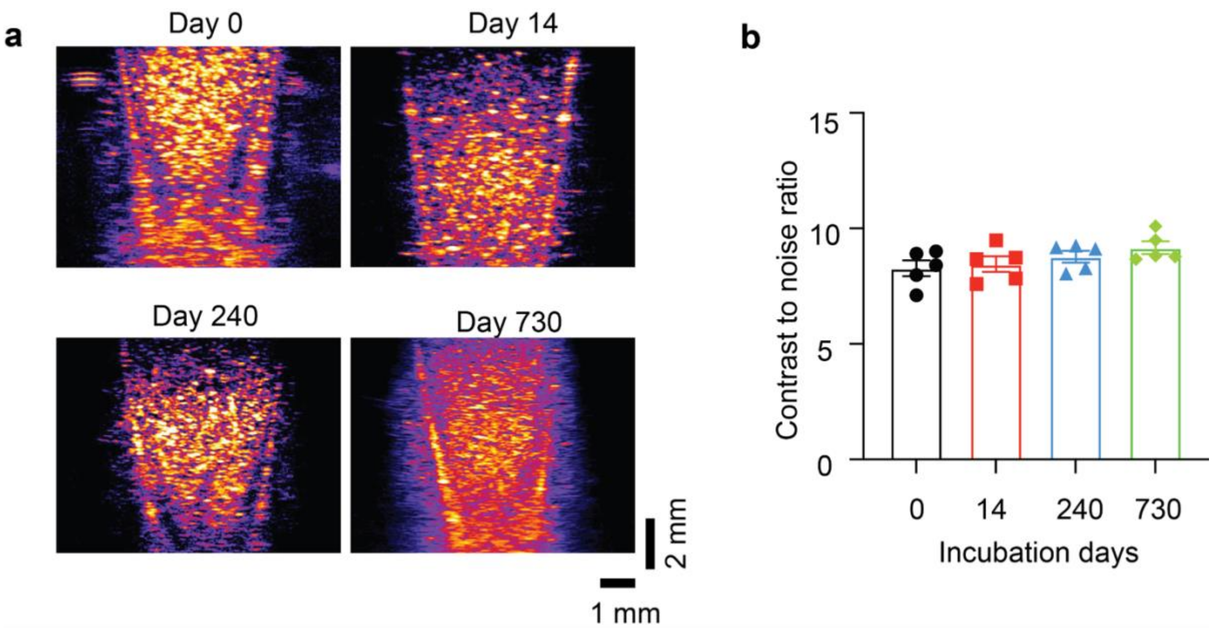


**Supplementary Fig. 28. Long-term ultrasound imaging stability of ZIF-8 liquid-phase gas nanotransducers at room temperature.** (a) Ultrasound images of silicone phantoms containing ZIF-8 liquid-phase gas nanotransducers acquired at 57 MHz after storage at room temperature for the indicated durations, up to 2 years. (b) Quantification of the contrast-to-noise ratio (CNR) of ZIF-8 liquid-phase gas nanotransducers following long-term storage at room temperature for up to 2 years.

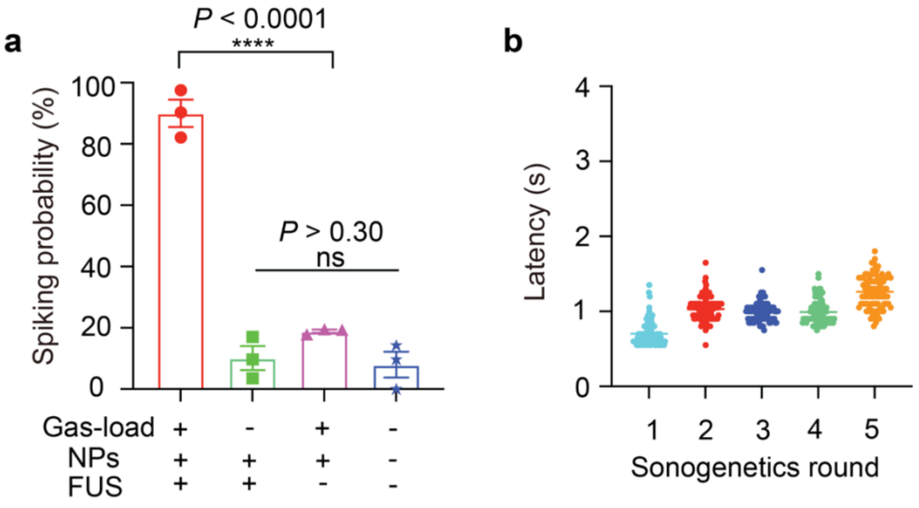


**Supplementary Fig. 29. Quantification of neuronal spiking under ultrasound stimulation.** (a) Spiking probability of primary neurons under different conditions (n = 100 neurons per group, from three independent experiments). Data are presented as mean ± s.e.m. Statistical analysis by two-way ANOVA with Tukey's multiple comparisons test (ns, $P \geq 0.05$; ****, $P < 0.0001$). (b) Spiking latency of primary neurons under sonogenetic stimulation (n = 100 neurons per group, from three independent experiments). Data are presented as mean ± s.e.m.

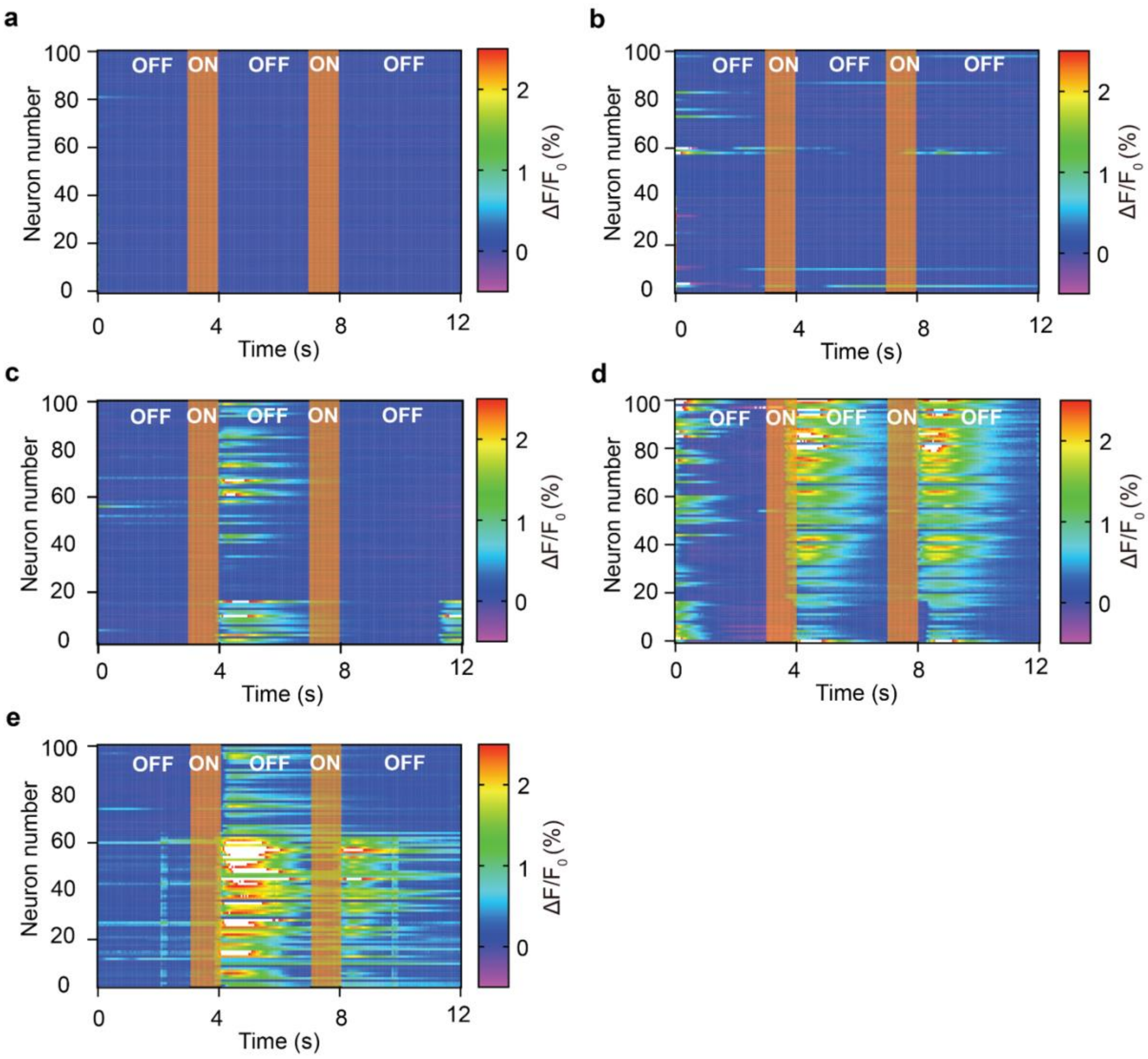


**Supplementary Fig. 30. Pressure-dependent neuronal activation under ultrasound stimulation.** Heat maps of normalized GCaMP6s fluorescence intensity from primary neurons treated with gas-loaded nanotransducers under ultrasound stimulation at different peak pressures: (a) 0 MPa, (b) 0.04 MPa, (c) 0.11 MPa, (d) 0.19 MPa and (e) 0.34 MPa (n = 100 neurons per group, from three independent experiments).

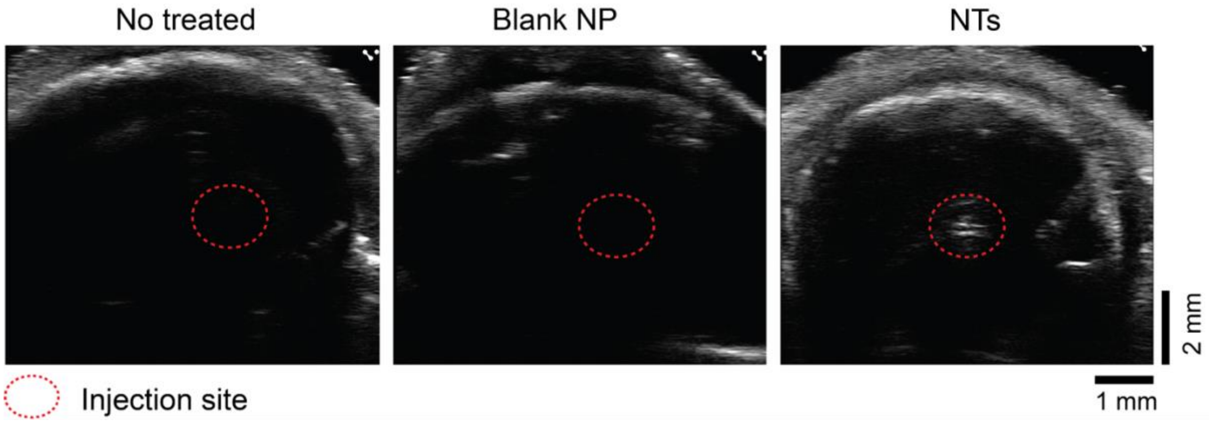


**Supplementary Fig. 31.** Ultrasound imaging of non-injected controls, control nanoparticles (gas-free ZIF-8), and gas-loaded ZIF-8 nanotransducers following injection into the VTA.

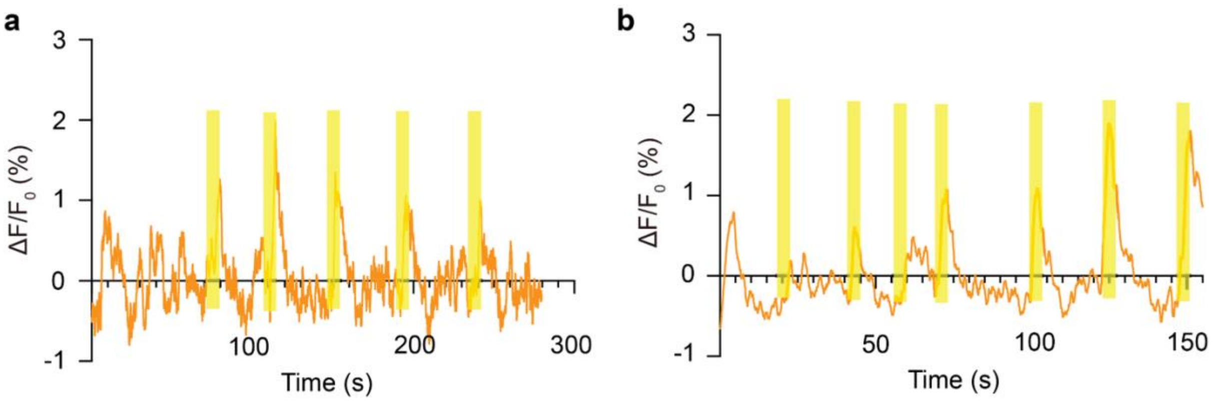


**Supplementary Fig. 32.** Normalized GCaMP6s fluorescence changes ($\Delta F/F_0$) in the mouse VTA under sonogenetic stimulation at (a) 4 weeks and (b) 12 weeks post-injection. FUS parameters: 1.5 MHz, 0.55 MPa, 3000 ms stimulation duration, 300 ms pulse width, 40% duty cycle, and 1 kHz PRF.

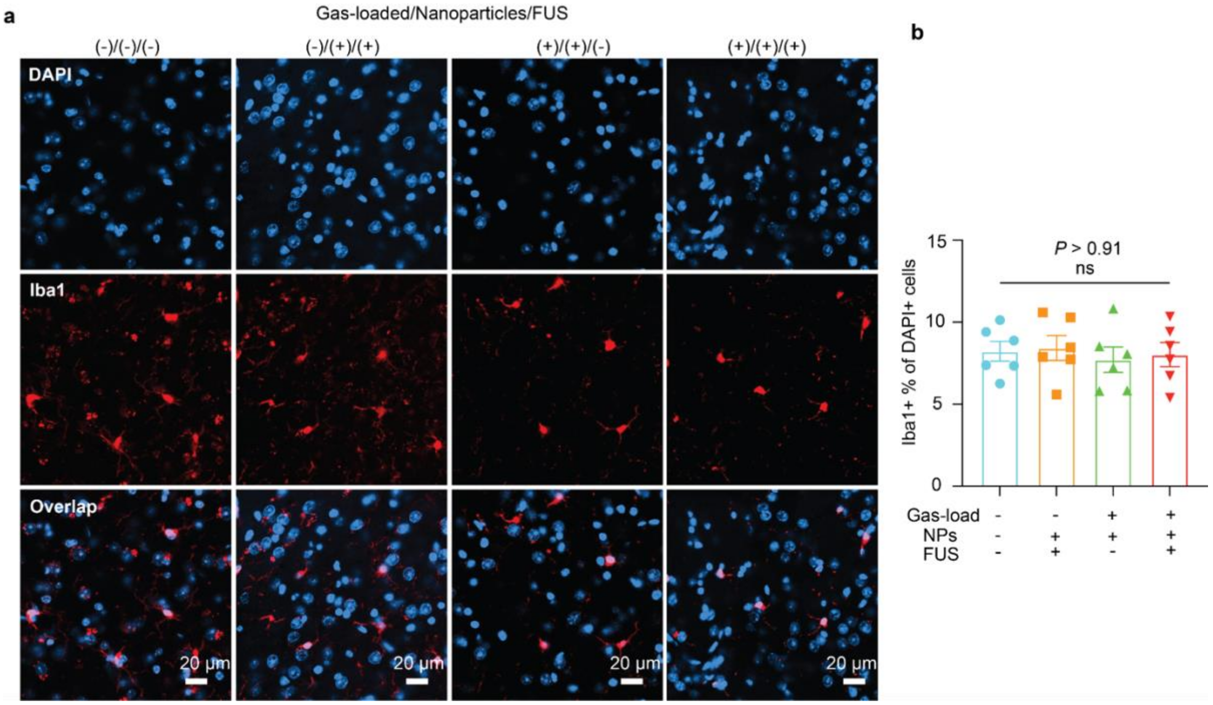


**Supplementary Fig. 33. *In vivo* biocompatibility evaluation of the sonogenetics via determining microglia (Iba1) activation.** (a) Fluorescence images of Iba1 of the brain slices in mouse VTA under the different conditions 14 days after sonogenetic stimulation. Scale bar : 20 μm. The blue signal is DAPI, and the red signal is Iba1. (b) Statistical analysis of the Iba1 intensity. Mean ± SEM, n =6, mice in each group. Two-way ANOVA and Tukey's multiple comparison test ($P \geq 0.05$ (ns), * $0.01 \leq P < 0.05$, ** $0.001 \leq P < 0.01$, **** $P < 0.0001$).

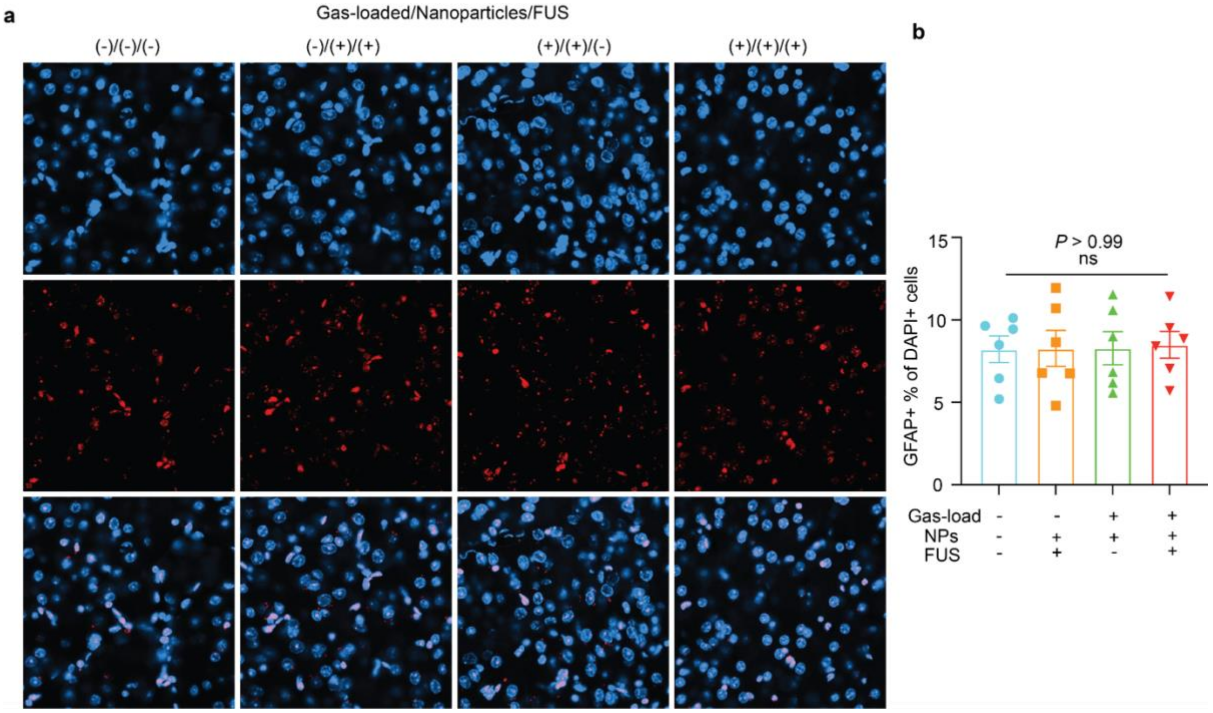


**Supplementary Fig. 34. *In vivo* biocompatibility evaluation of the sonogenetics via determining astrocytes (GFAP) activation.** (a) Fluorescence images of GFAP of the brain slices in mouse VTA under the different conditions 14 days after sonogenetic stimulation. Scale bar : 20 μm. The blue signal is DAPI, and the red signal is GFAP. (b) Statistical analysis of the GFAP intensity. Mean ± SEM, n =6, mice in each group. Two-way ANOVA and Tukey's multiple comparison test ($P \geq 0.05$ (ns), * $0.01 \leq P < 0.05$, ** $0.001 \leq P < 0.01$, **** $P < 0.0001$).

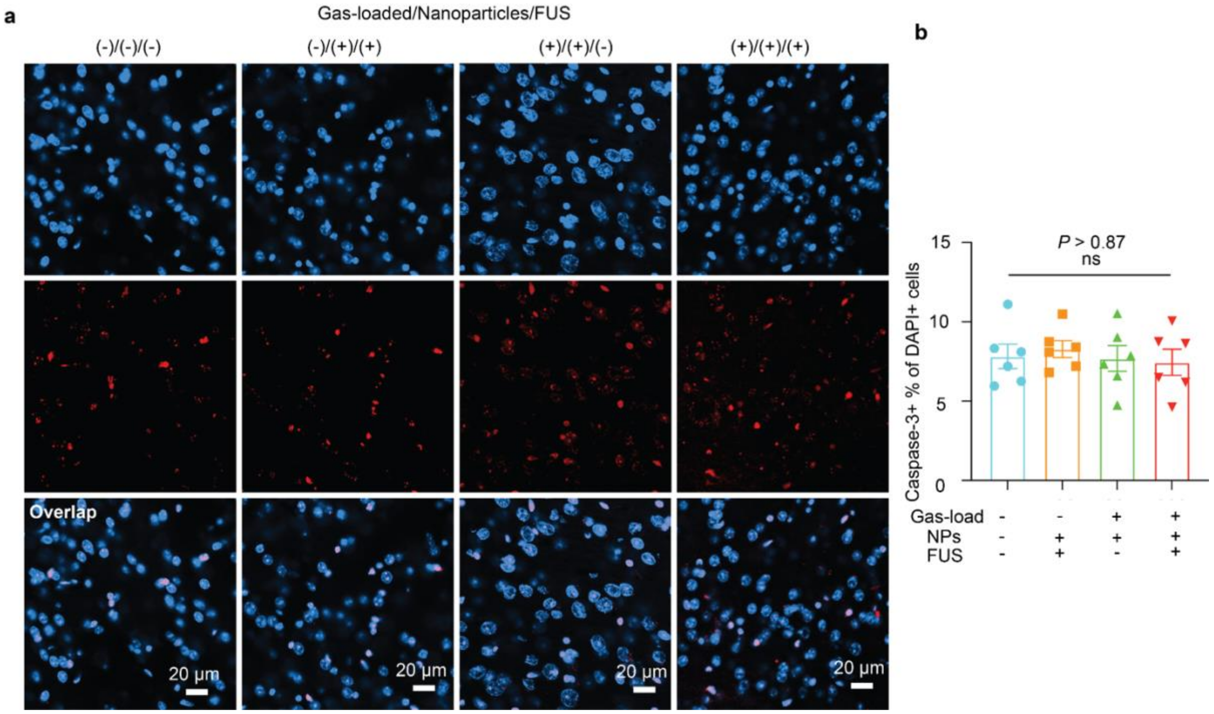


**Supplementary Fig. 35. *In vivo* biocompatibility evaluation of the sonogenetics via determining neuron apoptosis (Caspase-3).** (a) Fluorescence images of Caspase-3 of the brain slices in mouse VTA under the different conditions 14 days after sonogenetic stimulation. Scale bar : 20 μm. The blue signal is DAPI, and the red signal is Caspase-3. (b) Statistical analysis of the Caspase-3 intensity. Mean ± SEM, n =6, mice in each group. Two-way ANOVA and Tukey's multiple comparison test ($P \geq 0.05$ (ns), * $0.01 \leq P < 0.05$, ** $0.001 \leq P < 0.01$, **** $P < 0.0001$).

**Supplementary Table 1. Antibodies used in this work.**

| **Primary antibodies** | **Secondary antibodies** |
|---|---|
| Rabbit anti-Iba1<br>(1:500, 013-27691, Wako Chemicals) | Donkey anti-Rabbit, Alexa Fluor 594<br>(1:500, A32754, Invitrogen) |
| Rabbit anti-Cleaved Caspase-3<br>(1:500, 9661, Cell Signaling Tec.) | Donkey anti-Rabbit, Alexa Fluor 594<br>(1:500, A32754, Invitrogen) |
| Rabbit anti-GFAP<br>(1:500, 13-0300, Invitrogen) | Donkey anti-Rabbit, Alexa Fluor 594<br>(1:500, A32754, Invitrogen) |
| Mouse anti-tyrosine hydroxylase antibody (1:1000, MA1-24654, Fisher Scientific,) | Goat anti-mouse Alexa Fluor 488 (1:1000, ab150113, Abcam) |
| Rabbit anti-c-Fos antibody for mice<br>(1:500, ab222699, Abcam) | Goat anti-rabbit Alexa Fluor 405 (1:500, ab175652, Abcam) |
| | Hoechst 33342<br>(1:5000, 17535, AAT Bioquest ) |

**References**


1. D. P. Erdosy, M. B. Wenny, J. Cho, C. DelRe, M. V. Walter, F. Jiménez-Ángeles, B. Qiao, R. Sanchez, Y. Peng, B. D. Polizzotti, M. O. de la Cruz, J. A. Mason, Microporous water with high gas solubilities. *Nature* **608**, 712–718 (2022).

2. G. Ortiz, H. Nouali, C. Marichal, G. Chaplais, J. Patarin, Energetic performances of the metal-organic framework ZIF-8 obtained using high pressure water intrusion-extrusion experiments. *Phys. Chem. Chem. Phys.* **15**, 4888–4891 (2013).

3. G. Fraux, F.-X. Coudert, A. Boutin, A. H. Fuchs, Forced intrusion of water and aqueous solutions in microporous materials: from fundamental thermodynamics to energy storage devices. *Chem Soc Rev* **46**, 7421–7437 (2017).

4. D. Neumann, D. Woermann, Stability of the volume of air trapped on the abdomen of the water spider Argyroneta aquatica. *Springerplus* **2**, 694 (2013).

5. A. McGhee, A. Bennett, P. Ifju, G. W. Sawyer, T. E. Angelini, Full-field deformation measurements in liquid-like-solid granular microgel using digital image correlation. *Exp. Mech.* **58**, 137–149 (2018).

6. A. J. McGhee, E. O. McGhee, J. E. Famiglietti, K. D. Schulze, Dynamic subsurface deformation and strain of soft hydrogel interfaces using an embedded speckle pattern with 2D digital image correlation. *Exp. Mech.* **61**, 1017–1027 (2021).

7. J. Yang, H. C. Cramer III, C. Franck, Extracting non-linear viscoelastic material properties from violently-collapsing cavitation bubbles. *Extreme Mech. Lett.* **39**, 100839 (2020).

8. J. Yang, H. C. Cramer III, E. C. Bremer, S. Buyukozturk, Y. Yin, C. Franck, Mechanical characterization of agarose hydrogels and their inherent dynamic instabilities at ballistic to ultra-high strain-rates via inertial microcavitation. *Extreme Mech. Lett.* **51**, 101572 (2022).

9. J. Noack, A. Vogel, Laser-induced plasma formation in water at nanosecond to femtosecond time scales: calculation of thresholds, absorption coefficients, and energy density. *IEEE J. Quantum Electron.* **35**, 1156–1167 (1999).

10. X.-X. Liang, N. Linz, S. Freidank, G. Paltauf, A. Vogel, Comprehensive analysis of spherical bubble oscillations and shock wave emission in laser-induced cavitation. *J. Fluid Mech.* **940** (2022).

11. A. Vogel, S. Busch, U. Parlitz, Shock wave emission and cavitation bubble generation by picosecond and nanosecond optical breakdown in water. *J. Acoust. Soc. Am.* **100**, 148–165 (1996).

12. E.-A. Brujan, A. Vogel, Stress wave emission and cavitation bubble dynamics by nanosecond optical breakdown in a tissue phantom. *J. Fluid Mech.* **558**, 281 (2006).

13. J. B. Estrada, C. Barajas, D. L. Henann, E. Johnsen, C. Franck, High strain-rate soft material characterization via inertial cavitation. *J. Mech. Phys. Solids* **112**, 291–317 (2018).

14. D. Ranjan, J. Oakley, R. Bonazza, Shock-bubble interactions. *Annu. Rev. Fluid Mech.* **43**, 117–140 (2011).

15. M. Adama Maiga, O. Coutier-Delgosha, D. Buisine, A new cavitation model based on bubble-bubble interactions. *Phys. Fluids (1994)* **30**, 123301 (2018).

16. N. A. Pelekasis, A. Gaki, A. Doinikov, J. A. Tsamopoulos, Secondary Bjerknes forces between two bubbles and the phenomenon of acoustic streamers. *J. Fluid Mech.* **500**, 313–347 (2004).

17. J. Yang, V. Rubino, Z. Ma, J. Tao, Y. Yin, A. McGhee, W. Pan, C. Franck, SpatioTemporally adaptive quadtree mesh (STAQ) digital image correlation for resolving large deformations around complex geometries and discontinuities. *Exp. Mech.* **62**, 1191–1215 (2022).

18. J. Leu, Z. Tong, A. Doty, S. Tsimpoukis, B. Deng, J. Yang, Machine learning-aided spatial adaptation for improved Digital image correlation analysis of complex geometries. *Strain* **62** (2026).

19. A. McGhee, J. Yang, E. C. Bremer, Z. Xu, H. C. Cramer III, J. B. Estrada, D. L. Henann, C. Franck, High-speed, full-field deformation measurements near inertial microcavitation bubbles inside viscoelastic hydrogels. *Exp. Mech.* **63**, 63–78 (2023).

20. F. R. Gilmore, California Institute of Technology. Hydrodynamics Laboratory, *The Growth Or Collapse of a Spherical Bubble in a Viscous Compressible Liquid* (1952).

21. W. Wang, Y. Shi, W. Chai, K. W. K. Tang, I. Pyatnitskiy, Y. Xie, X. Liu, W. He, J. Jeong, J.-C. Hsieh, A. R. Lozano, B. Artman, X. Shi, N. Hoefer, B. Shrestha, N. B. Stern, W. Zhou, D. W. McComb, T. Porter, G. Henkelman, B. Chen, H. Wang, H-bonded organic frameworks as ultrasound-programmable delivery platform. *Nature* **638**, 401–410 (2025).

22. P. Li, S.-Y. Moon, M. A. Guelta, L. Lin, D. A. Gómez-Gualdrón, R. Q. Snurr, S. P. Harvey, J. T. Hupp, O. K. Farha, Nanosizing a metal-organic framework enzyme carrier for accelerating nerve agent hydrolysis. *ACS Nano* **10**, 9174–9182 (2016).
